# Global genomics of *Aedes aegypti* unveils widespread and novel infectious viruses capable of triggering a small RNA response

**DOI:** 10.1101/2024.06.06.597482

**Authors:** Shruti Gupta, Rohit Sharma, Adeline E. Williams, Irma Sanchez-Vargas, Noah H. Rose, Chao Zhang, Alexander Crosbie-Villaseca, Zheng Zhu, Gargi Dayama, Andrea Gloria-Soria, Doug E. Brackney, Jessica Manning, Sarah S. Wheeler, Angela Caranci, Trinidad Reyes, Massamba Sylla, Athanase Badolo, Jewelna Akorli, Ogechukwu B. Aribodor, Diego Ayala, Wei-Liang Liu, Chun-Hong Chen, Chalmers Vasquez, Cassandra Gonzalez Acosta, Alongkot Ponlawat, Tereza Magalhaes, Brendan Carter, Dawn Wesson, Darred Surin, Meg A. Younger, Andre Luis Costa-da-Silva, Matthew DeGennaro, Alexander Bergman, Louis Lambrechts, Carolyn S. McBride, Ken E. Olson, Eric Calvo, Nelson C. Lau

**Author notes:** Equal significant contribution (Shruti Gupta, Rohit Sharma).

## Abstract

The mosquito *Aedes aegypti* is a prominent vector for arboviruses, but the breadth of mosquito viruses that infects this specie is not fully understood. In the broadest global survey to date of over 200 *Ae. aegypti* small RNA samples, we detected viral small interfering RNAs (siRNAs) and Piwi interacting RNAs (piRNAs) arising from mosquito viruses. We confirmed that most academic laboratory colonies of *Ae. aegypti* lack persisting viruses, yet two commercial strains were infected by a novel tombus-like virus. *Ae. aegypti* from North to South American locations were also teeming with multiple insect viruses, with Anphevirus and a bunyavirus displaying geographical boundaries from the viral small RNA patterns. Asian *Ae. aegypti* small RNA patterns indicate infections by similar mosquito viruses from the Americas and reveal the first wild example of dengue virus infection generating viral small RNAs. African *Ae. aegypti* also contained various viral small RNAs including novel viruses only found in these African substrains. Intriguingly, viral long RNA patterns can differ from small RNA patterns, indicative of viral transcripts evading the mosquitoes’ RNA interference (RNAi) machinery. To determine whether the viruses we discovered via small RNA sequencing were replicating and transmissible, we infected C6/36 and Aag2 cells with *Ae. aegypti* homogenates. Through blind passaging, we generated cell lines stably infected by these mosquito viruses which then generated abundant viral siRNAs and piRNAs that resemble the native mosquito viral small RNA patterns. This mosquito small RNA genomics approach augments surveillance approaches for emerging infectious diseases.

## INTRODUCTION

The yellow fever mosquito, *Aedes aegypti*, has a global reach across most major human locales, and to this day remains a major health scourge as a prominent vector for arthropod-borne viral (arboviral) diseases like dengue fever. Municipal vector control organizations conduct routine molecular surveillance of arboviruses from trapped *Ae. aegypti* mosquitoes but are only able to assay known viruses^1^. Recent research efforts applying high-throughput RNA sequencing have now led to an explosion in the mosquito virome lists^2–5^. However, the persistence of these novel insect viruses within mosquito populations and their vertical transmission has not been fully examined in detail.

Vertical transmission and infection rates of arboviruses in *Ae. aegypti* have been examined for dengue and Zika viruses, among others^6^. Even when infection rates are high, the vertical transmission rates can be low if virus infection does not extend to the gonads^7–10^. Vertical transmission of arboviruses like Zika virus in laboratory *Ae. aegypti* infections is possible, but at low rates^11,12^. In addition, tracking prevalence of virus infections in mosquitoes in the wild is a complex epidemiological challenge when there are fluctuating levels of virus infection rates even during an outbreak^13–15^.

The ability of a virus to persist via vertical transmission in mosquito populations raises the concern of a species jump to humans^16,17^. Yet, tracking persistent viruses in wild and small pools of mosquitoes can be difficult with rapid surveillance screening because RT-PCR can be too imprecise to distinguish between strong viral RNA expression and low viral loads. Therefore, another molecular measure is needed to provide a better representation of persistent virus infections in mosquito populations.

Mosquitoes and mosquito cell cultures naturally mount a robust innate immune response to viruses with the RNA-interference (RNAi) pathway. During virus replication, viral double-stranded RNAs are processed into viral small interfering RNAs (siRNAs, ∼18-23 nucleotides long) that are loaded into Argonaute (AGO) proteins^18^. A second RNAi pathway that mainly regulates Transposable Elements (TEs) involves the Piwi proteins that bind Piwi-interacting RNAs (piRNAs, ∼24-35nt long), which have sequences complementary to TEs^19^. Since many TEs are evolutionarily related to viruses, mosquito Piwi proteins may generate viral piRNAs similar to how TE piRNAs are made. The viral siRNAs that are antisense to the viral mRNA will trigger AGO cleavage of viral mRNAs. While viral piRNAs may have antiviral properties in mosquitoes^20,21^, this response may be complicated by possible virus-to-virus interactions^22^.

When we first developed a Mosquito Small RNA Genomics (MSRG) pipeline^23^, we detected widespread insect virus persistence in mosquito cell cultures and in a few laboratory mosquitoes. A recent mosquito small RNA sequencing survey by the Marques lab profiled *Ae. aegypti* and *Ae. albopictus* samples from South America, Asia and Africa locales^22^. This study presented an emerging collection of insect viruses generating a viral small RNA response, potentially suggesting virus persistence in *Aedes* mosquitoes. However, this study then focused on two insect viruses, Humaita-Tubiacanga virus (HTV) and the bunyavirus Phasi-Charoen-Like virus (PCLV), and their role in potentially influencing dengue virus (DENV) competence^22^. The question then remains: How prevalent globally are *Ae. aegypti* mosquito viruses that trigger a small RNA response?

Here, we present the most comprehensive global survey of *Ae. aegypti* mosquito small RNAs to date, comprised of new laboratory strains and wild-caught and lab colonies from the Americas, Asia, and Africa continents. Our results significantly expand the catalog of insect viruses capable of propogating in mosquitoes and generating a small RNA response. We demonstrate experimental evidence for the transmissibility of these novel insect viruses to mosquito cell cultures to generate an RNAi response. Because these mosquito viruses can possibly develop tropism for mammalian cells, our small RNA genomics approach is a valuable addition to current molecular surveillance efforts to detect newly emerging pathogens vectored by mosquitoes and other arthropods.

## RESULTS

The collection of *Ae. aegypti* mosquitoes was organized through collaborators shipping mosquito samples that were either preserved (frozen or submerged in an RNA-protectant solution) or shipped as live animals for total RNA extraction and small RNA library preparation. Most mosquito RNAs were analyzed from whole mosquitoes to optimize for throughput.

Published small RNA libraries from Olmo et al^22^ and our previous study^23^ were integrated into our analysis of viral small RNA profiles. We compiled >280 small and long RNA sequencing datasets of *Ae. aegypti* from a select set of laboratory strains and from locations in North, Central and South America, Asia, and Africa (**Figure 1**). All datasets were analyzed by our MSRG pipeline, but this study primarily focused on Insect Specific Viruses (ISVs) and arboviruses while TEs and piRNA cluster loci analyses will be presented in a future study.

**Figure 1.**
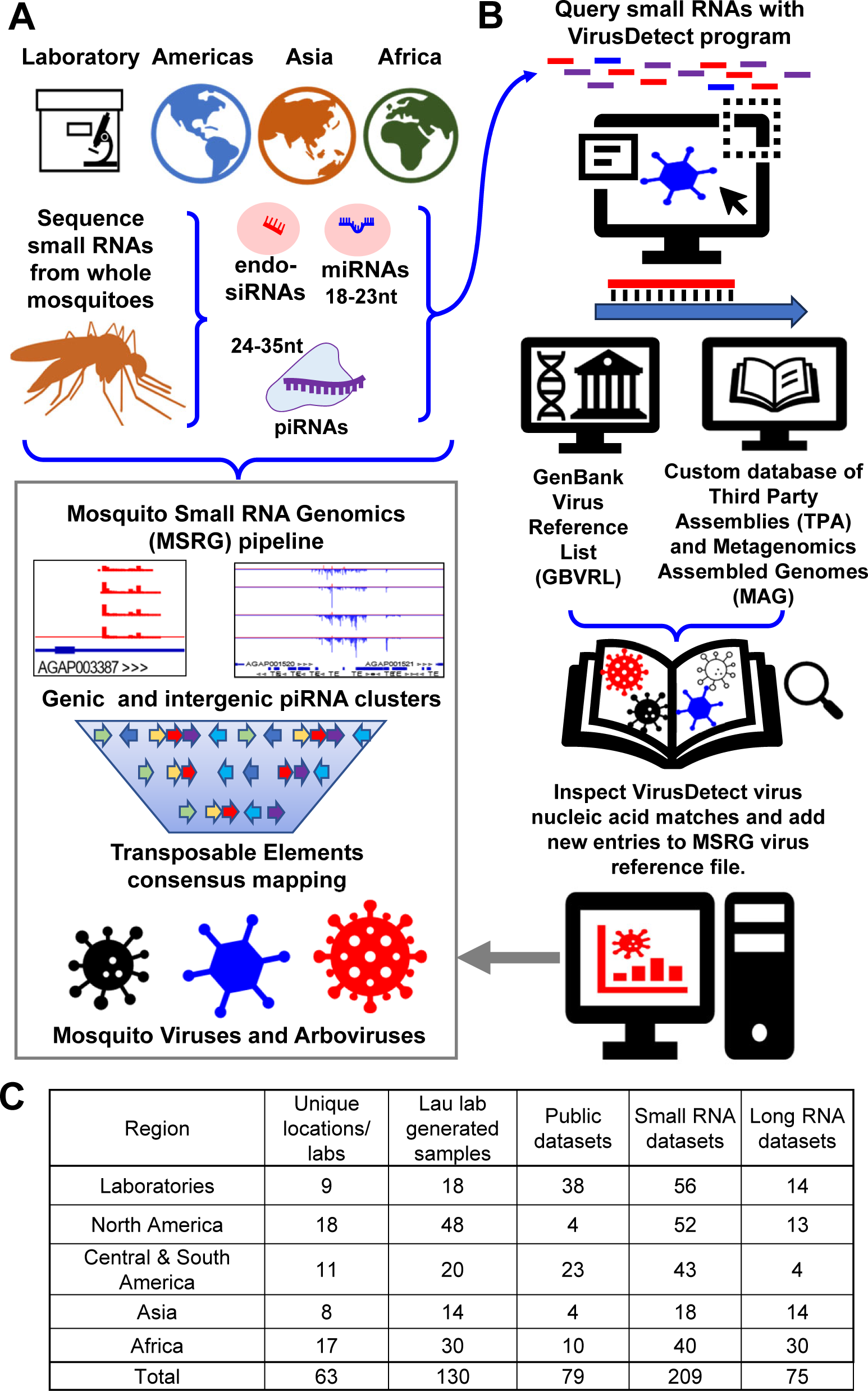
Overview of the global mosquito small RNA survey to discover RNA-interference (RNAi) responses to mosquito viruses. (A) Overview of the MSRG pipeline applied to a global survey of whole mosquitoes from Americas, Asia, Africa and laboratory strains. (B) Implementation of the VirusDetect program with updated GBVRL and custom databases for comprehensive mosquito virus detection. (C) Summary tabulation of the samples and RNA libraries analyzed in this study.

To address library quality control (QC), we first only selected libraries for analysis with at least 1 million reads. There was an average depth of ∼40M reads across all libraries (see **Supplemental Table S1 and S2** for library sequencing depth statistics). We also tracked an Endogenous Viral Element (EVE) in *Ae. aegypti* named AEFE1^24^, whose small RNAs are expressed in all mosquito samples we analyzed and are represented by primarily antisense piRNAs seen as blue peaks in coverage plots (**Supplemental Figure S1A**). Lastly, we inspected the read length distribution profiles for each library to track the expected peaks for miRNAs and piRNAs (**Supplemental Figure S2**). A virus needed to show at least 10 small RNA reads per million (RPM) to register on the bubble plots; if no viruses are detected, these QC measures confirmed a small RNA library was trustworthy.

### Augmenting the MSRG pipeline with improved virus discovery

Mosquitoes’ notable somatic piRNAs^25^ lend to the broad diversity of small RNA sequences in whole animal total RNA. This breadth means that small RNA libraries cannot be queried efficiently or specifically against databases such as the GenBank Virus Reference List (GBVRL) that have become flooded with recent massive virome surveys^4,5^ and coronavirus variant genomes^27^. In GBVRL version-249, there were ∼7.4 million records and 53.7% of these were beta-coronaviruses. To overcome this inefficiency and provide scalability, MSRG utilizes curated lists of viruses and TEs for small RNAs to be mapped against using BowTie (v1)^26^.

To address the limitations of the manually curated virus database in the MSRG, we devised an auxiliary pipeline to first analyze all *Ae. aegypti* small RNA libraries with the VirusDetect program^28^ (**Fig. 1B**). The VirusDetect program first performs *de novo* assembly of small RNAs into long contigs that can then query GBVRL efficiently and specifically to yield the count and coverage of the contigs against the most updated virus list in GBVRL (**Supp. Fig. S1B,C**). In this study, we updated the GBVRL in VirusDetect and locked it at version-249 from June 2022. The VirusDetect results confirmed the presence of viruses already present in the 2019 MSRG virus database while also revealing new viruses that we then added to our MSRG virus database list (**Supplemental Table S3**).

We then set up additional VirusDetect queries of our mosquito small RNA datasets against the Third-Party Assembly (TPA) and Metagenomics Assemble Genomes (MAG) databases that are distinct from GBVRL. This step was needed because when VirusDetect was run on GBVRL alone, a false-positive call was made for a tombus-like virus isolated from a *Hyposignathus monstrosus* bat^29^ due to the low contig coverage (**Supp. Fig. S1D,E**). The additional TPA and MAG runs discovered the Tiger Mosquito Bi-segmented Tombus-Like virus^30^ (TMBTLV) as the true source of these small RNAs, as well as revealing a new *Aedes* partiti-like virus^30^. In total, the VirusDetect analysis added 94 new viruses to our MSRG virus list (**Supp. Table S3**).

### Academic and commercial laboratory strains of Ae. aegypti

Limited insect viruses were found in academic lab strains from these new *Ae. aegypti* small RNA libraries as well as from previously examined lab strain datasets (**Figure 2A**). Our first MSRG pipeline study^23^ also revealed very few insect viruses in these strains, but the expanded virus list and low viral reads in this study boosted our confidence in the sparseness of viruses in these strains. Main figure bubble plots show all small RNAs (18-35nt), while breakdowns of the siRNAs (18-23nt) and piRNAs (24-35nt) are shown in **Supplemental Figure S3.**

**Figure 2.**
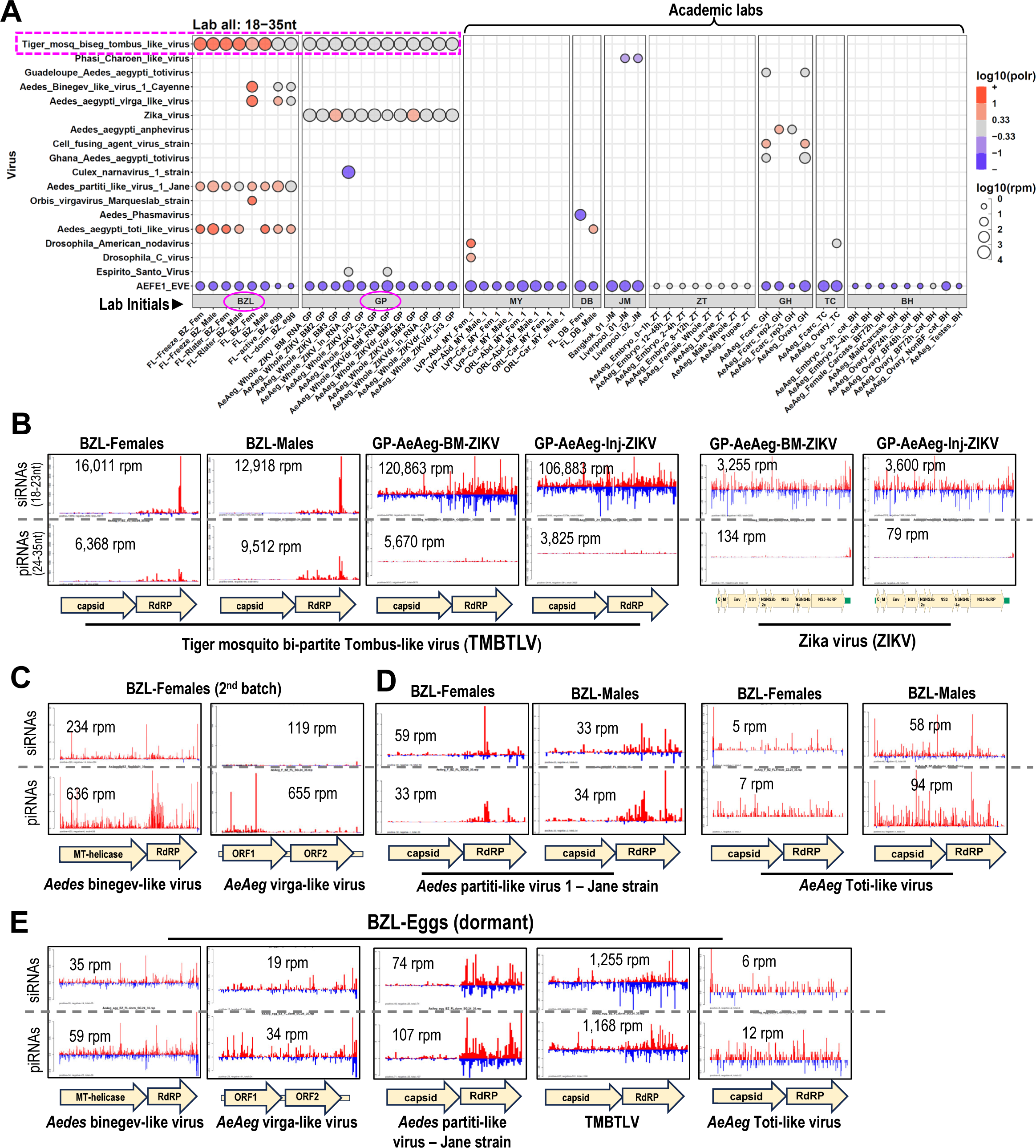
In contrast to commercial *Aedes aegypti* lab strains, most academic lab strains are low in viral small RNAs indicative of persistent viruses. (A) Bubble plot of viral small RNAs from lab strains. Number reads per million is reflected by bubble diameter, and color represents strand bias of reads, red is plus strand biased, blue is minus strand biased. Magenta circles and dashed boxes mark the BZL and GP samples that are commercial lab strains of *Ae. aegypti* and both are infected by Tiger mosquito bi-partite Tombus-like virus (TMBTLV). All the other *Ae. aegypti* samples were reared in academic labs. See metadata in Table S1 for sample details. Lab initials: BZL = Benzon Research, MY = Meg Younger, DB = Doug Brackney, JM = João Marques, ZT = Zhejian Tu, GH = Grant Hughes, TC = Tonya Colpitts, BH = Bruce Hay, GP = Gorben Pijlman (B) Coverage plots of TMBTLV and Zika virus (ZIKV) small RNAs from the two commercial lab strains. Additional coverage plots of (C) two viruses that only generated viral small RNAs in a 2^nd^ batch of female BZL *Ae. aegypti* and (D) other viruses always present in the BZL strain in both females and males. (E) Viral small RNAs and likely persistent viruses are detected in the BZL eggs at lower small RNA levels with a more distinct pattern of antisense viral small RNAs in eggs compared to the parental whole females in (B).

In contrast to these academic lab strains, two independent commercial lab strains from Benzon Research Labs (BZL) and Bayer AG (GP)^31^ contained massive amounts of TMBTLV small RNAs, with viral siRNAs more abundant than viral piRNAs (**Fig. 2B**). Coverage patterns of the TMBTLV small RNAs differed greatly between the two commercial strains. The BZL strain viral small RNAs were biased for the plus strand and mainly derived from the RNA dependent RNA Polymerase (RdRP) gene, while the GP strain displayed a much greater accumulation of minus strand siRNAs across the entire TMBTLV genome and had piRNAs still biased for the plus strand. All GP strain samples were co-infected with Zika virus (ZIKV), but control non-Zika-infected GP strains small RNA data were not available, so ZIKV’s contribution to the massive TMBTLV siRNAs cannot be resolved until future experiments can be conducted.

Commercial labs often raise *Ae. aegypti* alongside many other arthropods for the testing of pesticides. Benzon Research Labs states that their BZL strain was derived from an USDA “Gainesville” strain from 1994. The continuous rearing in their facility for over 25 years may have lent to this strain contracting additional viruses not seen in academic lab strains (**Fig. 2C,D**). These viruses must persist by vertical transmission, because the same viral small RNAs in adult BZL females were also detected in BZL eggs (**Fig. 2E**), although the eggs had lower levels and more minus strand viral small RNAs. For some mosquito viruses, the small RNAs originated from all genes in the viral genomes, but for a partiti-like virus and TMBTLV, the RdRP genes served as the primary source of small RNAs.

### Diverse profiles of viral small RNAs in Ae. aegypti across the two American continents

With the help of municipal vector control departments in California and Miami, Florida, along with collections and lab-maintained colonies from the DeGennaro, Olson, and Lambrechts labs, we assembled a diverse collection of *Ae. aegypti* samples from the American continents to generate and sequence small RNAs from whole mosquitoes. We also integrated previously published *Ae. aegypti* small RNA libraries from Suriname and Brazil^22^.

There was widespread prevalence of mosquito viruses generating robust viral small RNAs in North, Central and South America (**Figure 3A,B**). We detected other mosquito viruses related to plant viruses, similar to TMBTLV, from the two commercial lab strains (**Fig. 2B**). A Liverpool Tombus-like virus (LTLV) was also reported in the same metagenomics study discovering TMBTLV^30^, whereas Verdadero virus is a partitivirus first described infecting *Ae. aegypti* as well as *Drosophila*^32^. Partitiviruses were originally discovered to infect plants, protozoans, and fungi, while tombusviruses are a group representing the tomato bushy stunt viruses^33,34^. Viral siRNAs and piRNAs for LTLV were much higher in the midguts versus the total abdomens of an *Ae. aegypti* strain from Recife, Brazil (**Supplemental Figure S4A**), suggesting that mosquitoes feeding on plant sap nutrients could be a route for the transfer of plant-related viruses to an insect host.

**Figure 3.**
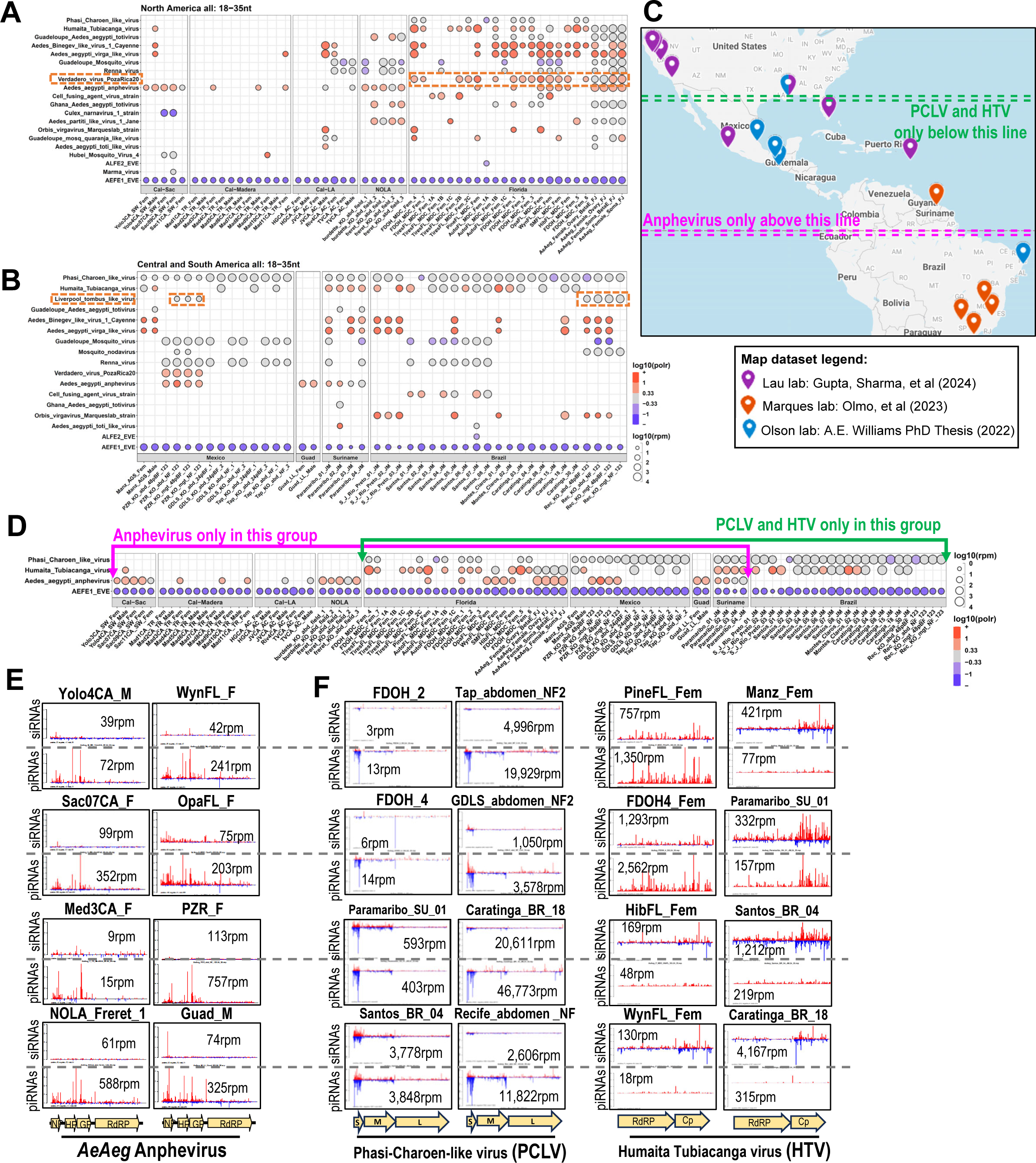
*Ae. aegypti* in the Americas display diverse insect virus small RNA responses with geographic boundaries delineated by three prevalent mosquito viruses. (A) Bubble plot of viral small RNAs from North America *Ae. aegypti* samples. Number reads per million is reflected by bubble diameter, and color represents strand bias of reads, red is plus strand biased, blue is minus strand biased. (B) Bubble plot of viral small RNAs from Central and South America *Ae. aegypti* samples. Dashed brown line boxes mark two viruses noted in Figure S4AB as being related to plant viruses. (C) Map of the Americas where the *Ae. aegypti* samples originated from, with geographic boundaries delineated by the distribution of Anphevirus and PCLV displayed in the bubble plot in (D). Example coverage plots of Anphevirus small RNAs (E) and PCLV and HTV small RNAs (F).

Even though many of the samples in these regions contained the same viruses, the coverage pattern of these viruses displayed remarkable differences. There were variations in the patterns of Verdadero viral small RNAs between a Mexico isolate and several different samples of Miami, Florida isolates (**Supp. Fig. S4B**). The Binegev-like virus and Renna virus were found from the west to the east coasts of the United States, and they also displayed interesting variations in viral small RNA patterns between samples (**Fig. 3A, Supp. Fig. S4D,E**).

The viruses most widespread across the *Ae. aegypti* small RNA samples of the Americas were PCLV and HTV (as noted previously in Olmo, et al^22^), along with the *Ae. aegypti* Anphevirus first described in a Florida strain^25^. Remarkably, PCLV and HTV were most frequently found in *Ae. aegypti* of the southern fraction of the Americas while Anphevirus displayed a bias for the northern fraction of the Americas (**Fig. 3C,D**). Florida, Mexico, and Caribbean locales reflected a mixing zone for these three viruses. Viral small RNA patterns and abundances fluctuated widely between individual mosquito isolates (**Fig. 3E,F**), but Anphevirus and HTV small RNAs had a clear plus strand bias, especially for the viral structural genes. In contrast, PCLV small RNAs displayed a stronger antisense bias and mostly originated from the ‘Small’ and ‘Medium’ segments. A principal component analysis of the viral small RNAs supports the delineation of North American group mosquitoes from the Central and South American mosquitoes (**Supp. Fig. S4F**).

Miami, Florida and various places in Brazil are sites of recent dengue fever outbreaks^35^, yet no DENV small RNAs were detected in any of these samples, including those from *Ae. aegypti* sampled by the Florida Department of Health (FDOH) in the vicinity of known dengue fever patients. The lack of DENV small RNAs mirrors the sporadic and challenging detection efforts of DENV in surveillance of mosquitoes to precede a human disease outbreak^1,13–15^. However, all of these samples revealed that multiple insect viruses can simultaneously infect and generate a robust RNAi response of viral small RNAs in a small mosquito cohort (**Fig. 3A, B**).

#### A snapshot into Asian Ae. aegypti viral small RNA patterns

We generated a new collection of small RNA libraries from *Ae. aegypti* mosquitoes across eastern Asia (**Figure 4A,B**) and observed a significant diversity of insect viruses generating significant small RNA responses. Several mosquito viruses in Asia were common with those in American samples (**Fig. 3**), such as HTV and PCLV (**Fig. 4C,D**). This also included the Partiti-like virus-1 Jane strain, which notably persisted in the ThaiKP colony that was propagated for >40 generations since its isolation in 2010^36^. Interestingly, the ThaiKP Partiti-like virus-1 viral small RNAs were strongly biased for the plus strand of the RdRP gene in contrast to both plus and minus strand siRNAs against this virus in a Singaporean wild isolate (**Fig. 4E**). The molecular implications of these and similar diverse fluctuations in viral small RNAs from HTV (**Fig. 4C**) remain unknown.

**Figure 4.**
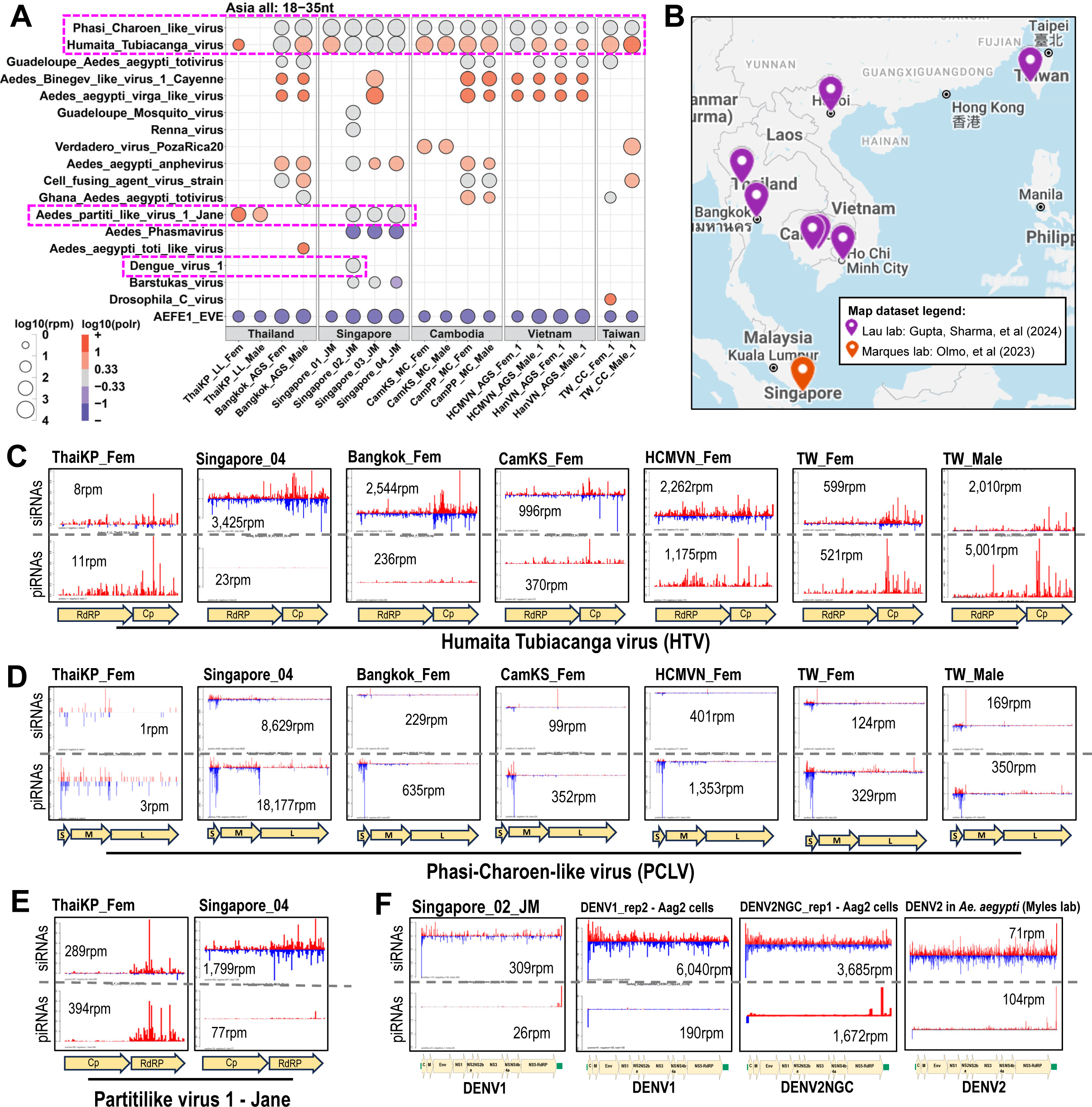
Asian *Ae. aegypti* viral small RNA patterns reveal a first case of dengue viral small RNAs from a wild isolate and share patterns with mosquitoes from the Americas. (A) Bubble plot of viral small RNAs from Asia *Ae. aegypti* samples. Number reads per million is reflected by bubble diameter, and color represents strand bias of reads, red is plus strand biased, blue is minus strand biased. Dashed pink line box mark notable viruses whose coverage plots are displayed below. (B) Map of Asia locations where the *Ae. aegypti* samples originated. Coverage plots of HTV (C), PCLV (D) and Partiti-like virus 1 Jane strain (E) viral small RNAs from a selection of Asian *Ae. aegypti*. (F) The DENV viral small RNA coverage from a Singapore isolate compared to other DENV viral small patterns from infections in Aag2 cells and a lab injected infection of *Ae. aegypti* by the Myles lab.

Surprisingly, our MSRG pipeline detected the first case of dengue viral small RNAs from a wild mosquito isolate – a Singaporean sample whose RNA was sequenced in Olmo, et al^22^ (**Fig. 4F**). DENV1 siRNAs in this Singaporean *Ae. aegypti* were more abundant than viral piRNAs, reflecting the same trend as DENV infections in mosquito Aag2 cells^23^. These natural DENV1 small RNAs also accumulated to the same level as a DENV2 infection of a lab *Ae. aegypti* experiment^37^, reflecting the likelihood of significant DENV1 infection in this Singaporean sample.

#### African Ae. aegypti colonies harbor a persistent African-specific mosquito virus

Most *Ae. aegypti* samples in this study thus far belong to the *Ae. aegypti aegypti* subspecies that have higher human biting proclivity and better ZIKV vector competence compared to the contemporary African subspecies *Ae. aegypti formosus*^38,39^. Most African colonies in this study are subspecies *formosus*, but some are subspecies *aegypti* (THI, NGO, CVerd), while others are substantially admixed (KUM, OGD). Distinct genetic backgrounds between *aegypti* and *formosus* subspecies are one likely explanation behind the differences in these subspecies^38^, but the persistent viruses generating viral small RNAs in *formosus* subspecies were previously unknown.

Our small RNA profiles revealed that PCLV’s global reach extends into several African colony strains primarily on the western African coast, which may be biased to the limited access to colonies only from these African locales (**Figure 5A,B**). The PCLV small RNA coverage patterns were also more biased towards the ‘S’ and ‘M’ segments with plentiful antisense viral piRNAs, while the ‘L’ segment mainly generated sense viral piRNAs (**Fig. 5C**). Unlike Asian and Central and Southern American *aegypti* strains where HTV frequently accompanied PCLV in causing a small RNA response in the same mosquito sample, *formosus* strains only had a few HTV small RNA signatures, all of which were found exclusive of PCLV (**Fig. 5A**).

**Figure 5.**
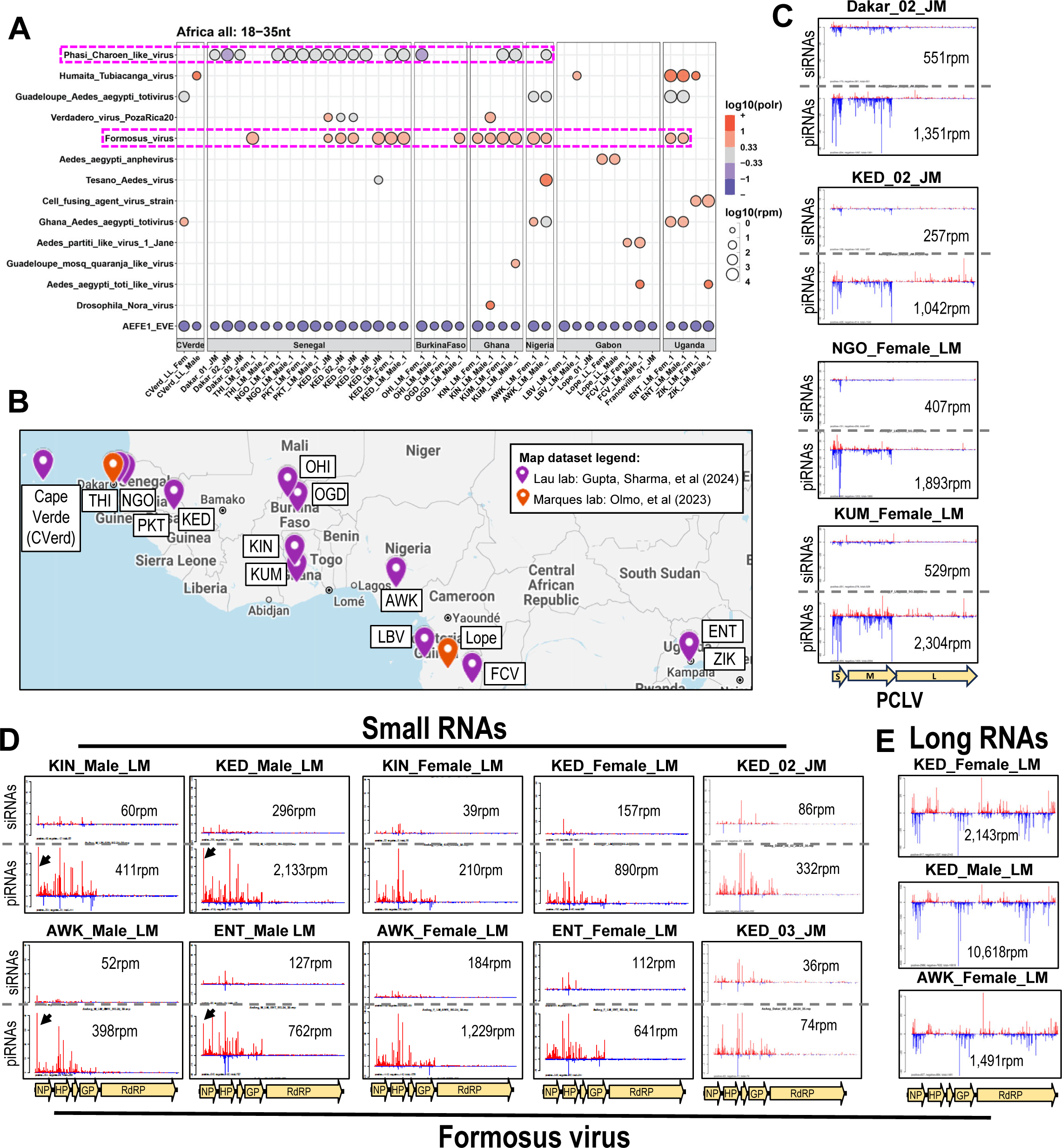
African *Ae. aegypti* colony strains carry viral small RNAs unique to their continent. (A) Bubble plot of viral small RNAs from Africa *Ae. aegypti* colony strains. Number reads per million is reflected by bubble diameter, and color represents strand bias of reads, red is plus strand biased, blue is minus strand biased. Dashed pink line boxes mark the PCLV and Formosus viruses noted in panels (C) and (D), respectively. (B) Map of Africa locations where the *Ae. aegypti* colonies or samples originated. (C) Coverage plots of PCLV small RNAs from a selection of African *Ae. aegypti* showing high M-fragment piRNAs rivaling the S-fragment piRNAs. (D) The Formosus virus viral small RNA coverage from African *Ae. aegypti* colonies from the McBride lab and an independent Kedougou, Senegal sample from Olmo et al 2023. The black arrow points to female-specific viral piRNA specie. (E) Three examples of Formosus virus long RNAs sequenced from matched samples in (D).

Formosus virus displayed the most striking persistent small RNA response in 12 samples from native-range colonies (**Fig. 5A**). This rhabdovirus has a ∼12.2kb genome that was initially absent from GBVRL database. It was deposited in the TPA (accession BK059424) by Parry, et al^30^, from a metagenomics assembly of transcriptomes from a different African lab colony not in this study originating from Bundibugyo, Uganda^40^. Importantly, four additional African *Ae. aegypti* samples from Olmo et al^22^, that were established and maintained separately from the main set of native-range colonies, also displayed Formosus virus small RNAs with similar coverage patterns (**Fig. 5D**, i.e. KED_02_JM and KED_Female_LM). Since most of the native-range colonies here were sampled after at least seven generations, we conclude that the Formosus virus is persisting and being transmitted vertically within the colonies.

Formosus virus primarily generates viral piRNAs from the sense strands of the NP, HP, and GP genes in the 5’ half of the viral genome, while the RdRP gene seemingly evades piRNA production (**Fig. 5D**). Conversely, long RNA sequencing indicated the plus-strand RdRP gene is transcribed as highly as NP, HP, and GP genes (**Fig. 5E**). We also detected robust minus strand long RNAs indicative of replication of the negative-strand rhabdovirus genome, yet much fewer viral piRNAs were generated from this minus strand (**Fig. 5D**). Future studies will investigate how the RdRP gene transcript and negative-strand genomic transcript evade the RNAi machinery. Lastly, we observed some sex-specific Formosus viral piRNA accumulation patterns in males at the 5’ end of the NP gene, even though long viral RNA patterns look similar between the sexes.

### Insights into mosquito virus replication and RNAi dynamics from matched long RNA sequencing

Next, we asked if the Formosus virus example of the interplay between viral small RNAs and long RNAs was indicative of a broader picture of dynamics between viral replication and viral small RNA biogenesis for other viruses in other *Ae. aegypti* mosquitoes. We compared in a scatterplot the log-transformed long and small RNA levels between ∼75 matched samples for each mosquito virus (**Figure 6A**). We noted four interesting virus groups in this analysis. The first group is AEFE1 EVE measurements from adults all clustering together, which displayed more small RNAs than long RNAs, and reflects the care we took to generate and sequence these RNA libraries as reproducibly as possible.

**Figure 6.**
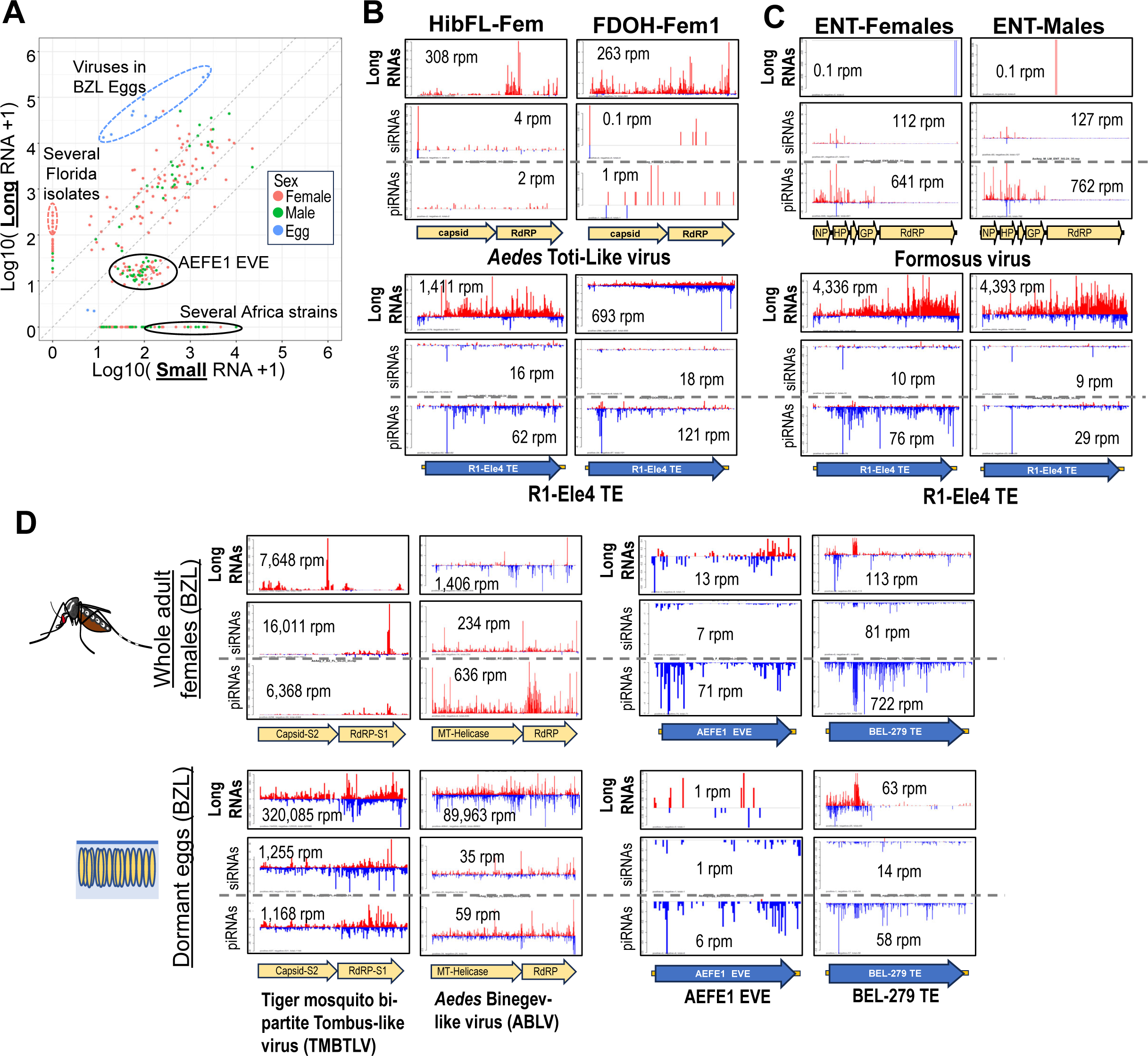
Virus transmission and replication dynamics suggested by long RNA sequencing compared to matched small RNAs of mosquito viruses and transposable elements (TEs). (A) Scatterplot comparing the particular *Ae. aegypti* samples that enabled matched library construction of small RNAs and long RNAs. Sequencing reads per million with a pseudo-count of 1 are plotted on the logarithmic scales. Dots representing the matched samples are colored by sex, and groups of samples clustering together are noted in the labeled ovals. (B) Coverage plots of two Florida isolates of *Ae. aegypti* exhibiting abundant long RNA signal for the totivirus but negligible viral small RNA in the upper plots that contrast both long and small RNAs against an *R1-Ele4* TE. (C) Coverage plots of long RNAs versus small RNAs for the Formosus virus and *R1-Ele4* TE from both males and females of the ENT African colony. (D) Coverage plots of long RNAs compared to small RNAs for viruses and a TE from the BZL strain of *Ae. aegypti* females and dormant eggs.

The next two groups were viruses in samples that displayed significant viral long RNAs but few small RNAs, and vice versa significant viral small RNAs without much long RNAs. In several Florida isolates that were captured by municipal vector control surveillance, the *Aedes* Toti-like virus appeared to replicate and express viral genes effectively, perhaps before the mosquitoes could mount an RNAi response with small RNAs (**Fig. 6B**). Notably, these Florida mosquitoes had robust small RNA responses to endogenous TEs like *R1-Ele4* as well as other viruses like Verdadero virus, validating that the lack of *Aedes* Toti-like virus small RNAs is not merely a technical error (**Supp. Fig. S4B**). More interestingly, there were some virus cases like Formosus virus in the Entebbe, Uganda colony (ENT), in which both males and females only generated viral small RNAs. This may be linked to the complete loss of the viral long RNAs (**Fig. 6C**) despite the long RNA libraries still tracking the TE long RNAs.

The fourth standout group is viruses in the BZL strain eggs which displayed massive levels of long RNA reads from both plus and minus strands of the various viruses in this strain (**Fig. 6D**). For both TMBTLV and an *Aedes* Binegev-like virus (ABLV), the long RNA reads were more than an order of magnitude greater than the small RNAs in both the eggs and whole adult females. Despite the production of abundant viral piRNAs and siRNAs in the adult whole female and maternal contribution of these viral small RNAs to the eggs, virus silencing does not appear effective, which could allow for efficient vertical virus transmission from female to egg.

### Infectious capacity of metagenomically assembled virus entries from RNA sequencing

Although TMBTLV and Formosus virus were in the TPA and MAG databases within GenBank^30^, they were not included in the GBVRL at the time of our initial analysis, raising the question of whether these entries are truly infectious viruses. To bolster our RNA sequencing findings, we sought to isolate viruses from filtered mosquito homogenates to infect mosquito cell cultures (**Figure 7A**). If the virus infections in cell cultures were deemed stable, we could then sequence the virus genomes for GBVRL submission and sequence viral small RNAs from infected cells to compare against the mosquito viral small RNAs.

**Figure 7.**
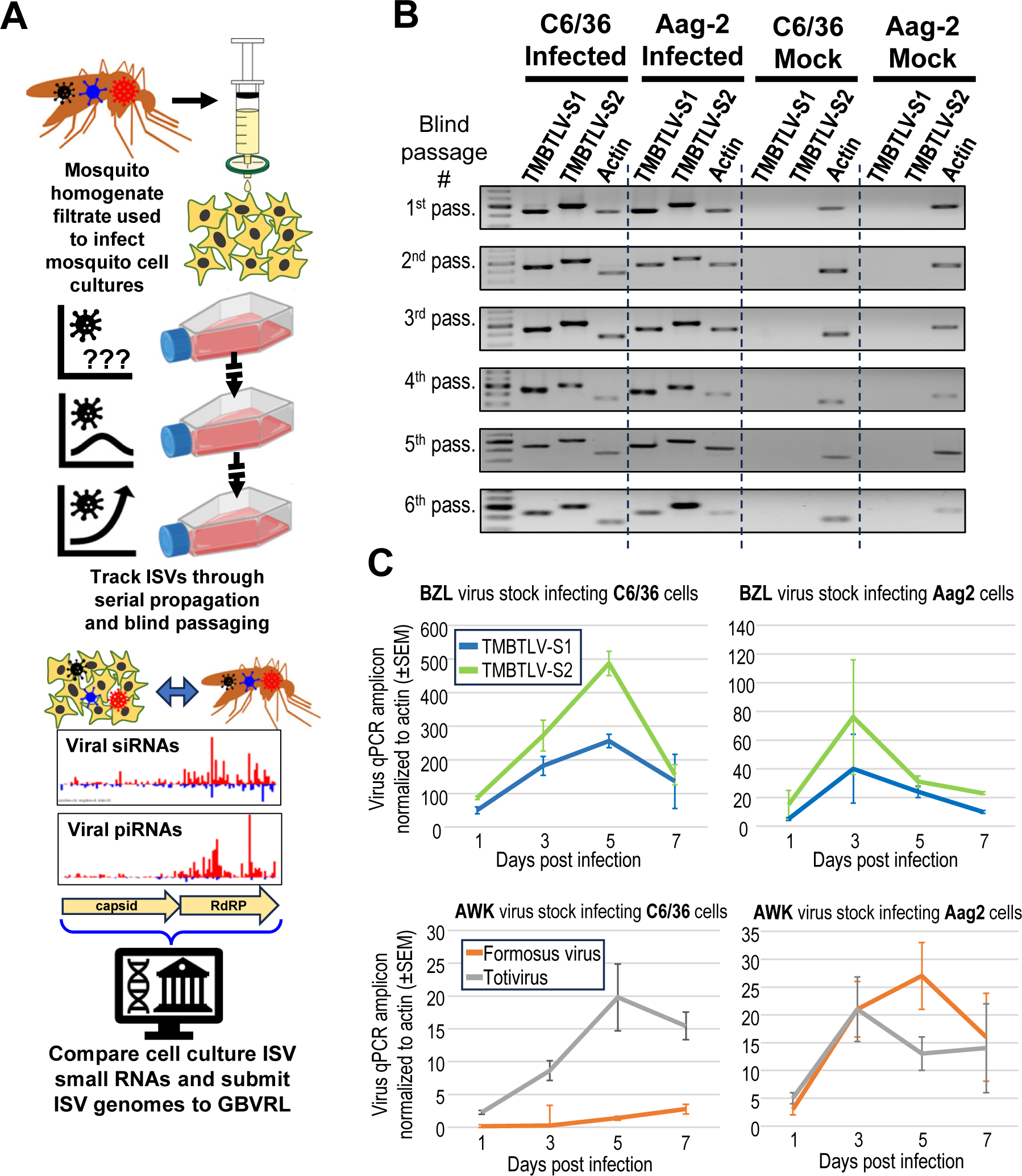
Novel mosquito viruses infecting and replicating in mosquito cells. (A) Our methodology to molecularly validate the small RNA detection of ISVs are true viruses that can be isolated and verified for triggering the RNAi response in mosquito cells. (B) RT-PCR detection of TMBTLV RNAs S1 and S2 during multiple rounds of blind passaging, starting with BZL mosquito homogenate as the initial virus infection source placed onto C6/36 and Aag2 mosquito cells. (C) Virus replication kinetics measured in C6/36 and Aag2 cells over 1 week. Flasks of 1 million cells were infected on Day 0 with 200 viral qPCR amplicons per infection. Virus stocks are from filtered media from subsequent passage from the experiment in (A). Error bars correspond to the standard error from measuring three biological replicates of infection of the mosquito cells with the same virus stock.

We followed established serial propagation and blind passaging procedures using C6/36 *Ae. albopictus* cells and Aag2 *Ae. aegypti* cells to test for virus infection from the homogenates of the BZL lab strain (**Fig. 7B**), a colony strain from Poza Rica, Mexico (PZR, **Supplemental Figure S5A**), and two African colony strains AWK and KIN (**Supp. Fig. S5B**). After 5-to-6 rounds of blind passaging, we were able to use RT-PCR and primers to consistently detect the stable infections of TMBTLV RNAs S1 and S2, the RdRP and Capsid genes of Verdadero virus, and a Totivirus and Formosus virus amplicon. These results suggested we could establish stably infected mosquito cells as new stocks of these mosquito viruses, albeit at moderate virus copy levels, since we have not yet established a titer regime for these viruses.

For TMBTLV, Totivirus, and Formosus virus, we examined infection kinetics in mosquito cells and virus tropism. TMBTLV infection rates proceeded faster and to a greater extent in C6/36 cells than Aag2 cells (**Fig. 7C**), likely because of the *Dcr2* mutation in C6/36 cells that reduces antiviral RNAi and makes C6/36 cells the most common cell culture system to isolate viruses from insects^41,42^. In contrast, Formosus virus exhibited efficient replication only in Aag2 cells, suggesting that this virus has more restricted species tropism than TMBTLV. Totivirus replicated in both cell types, although replication kinetics were a little faster in Aag2 cells. Lastly, we tested if these three viruses can also replicate in mammalian Vero E6 and Huh7.5 cells that are susceptible to mosquito-borne human viruses like ZIKV and DENV. Surprisingly, blind passage of BZL mosquito homogenates into these mammalian cells was effective at showing TMBTLV replication, and some cytopathic effects were observed (**Supp. Fig. S5C, D**). A fainter signal for the Formosus virus was also observed in some of the blind passaging in the mammalian cells.

The transfer of these mosquito viruses into cell cultures enabled us to clone TMBTLV and Formosus virus genomic fragments for sequencing and GBVRL submission. When comparing the genomes of our TMBTLV and Formosus virus isolates to the initial genomes assembled by Parry et al,^30^, we detected 25 total protein-coding mutations in the ∼4.5kb TMBTLV genome, but only 11 protein-coding mutations in the ∼12.2kb Formosus virus genome (**Supp. Figure S6).**

Together, these results provide strong evidence supporting the infectious capacity of these persistent mosquito viruses even though the source mosquitoes displayed an RNAi response with viral small RNAs.

### Small RNA responses in virus-infected mosquito cells are comparable to the whole mosquito

We subjected our virus-infected C6/36 cells and Aag2 cells to small RNA sequencing to determine whether new viral small RNA patterns in these cell cultures could recapitulate the patterns in the whole mosquito. These C6/36 and Aag2 cell lines were obtained from other mosquito laboratories and already had known viruses persisting and generating small RNAs^23,43–45^ (**Figure 8A,B**). We were able to reconfirm the presence of Cell Fusing Agent Virus (CFAV) and PCLV in both of our ‘mock’ C6/36 and Aag2 cells^23^, as well as discover Sobemo-like viruses and densoviruses in our C6/36 and Aag2 cells, respectively.

**Figure 8.**
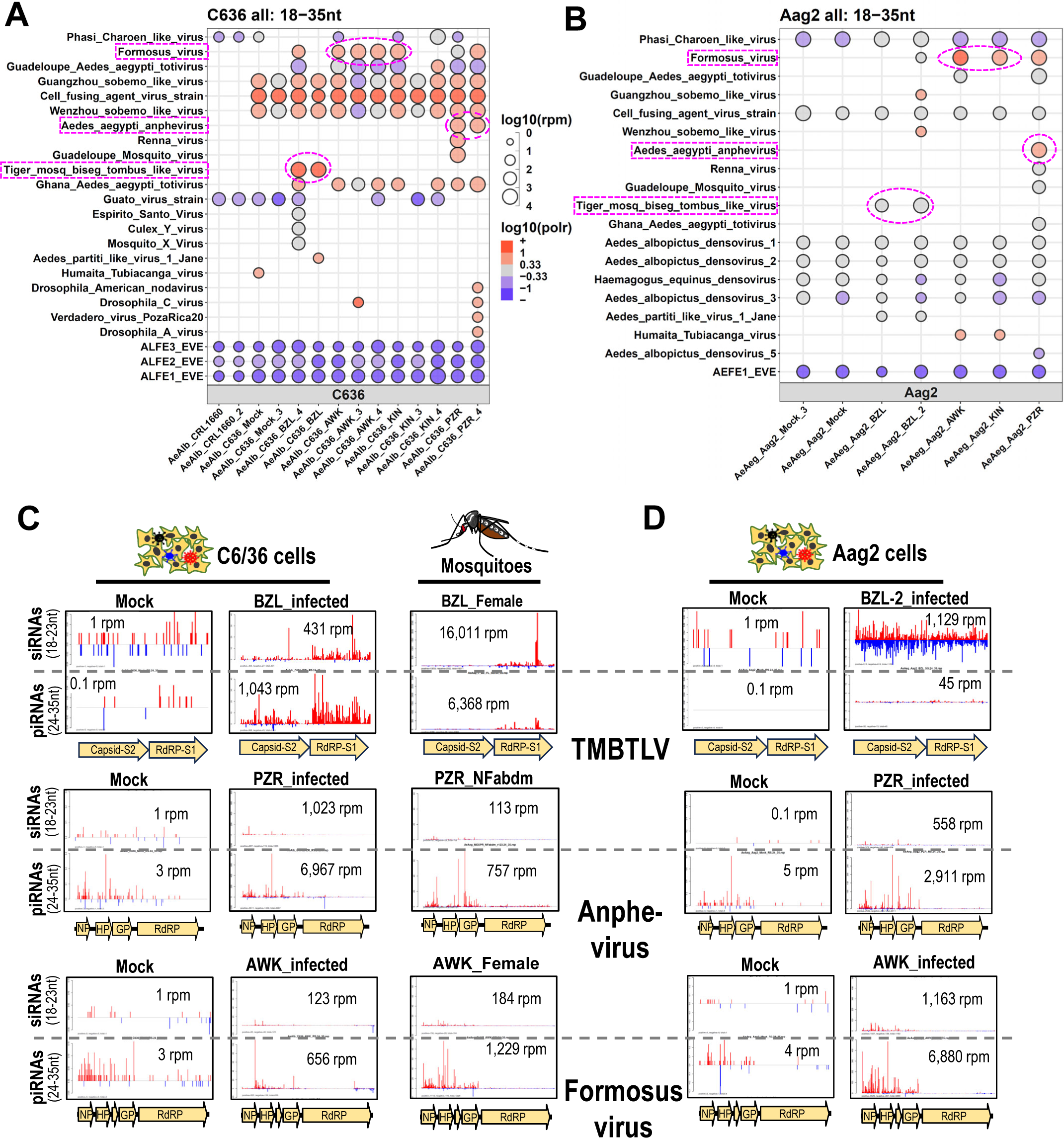
Mosquito viruses isolated in cell culture trigger an RNAi response of viral small RNA patterns that resemble the patterns from whole mosquitoes. Bubble plots of viral small RNAs sequenced from (A) *Ae. albopictus* C6/36 cells and (B) *Ae. aegypti* Aag2 cells, either mock or stably infected after multiple blind passages of virus stocks from mosquito homogenates. Number reads per million is reflected by bubble diameter, and color represents strand bias of reads, red is plus strand biased, blue is minus strand biased. Magenta boxes and dashed circles in (A) and (B) highlight specific samples inspected in coverage plots in (C) and (D). Coverage plots for three viruses, TMBTLV, Anphevirus and Formosus virus in (C) C6/36 cells and (D) Aag2 cells with the intact mosquito from females shown in the middle for comparison.

In addition to the preexisting viruses in the background-strain cell lines, these infected mosquito cells in Figures 7 and S5 exhibited viral small RNAs from TMBTLV (from BZL mosquitoes), Anphevirus (from PZR mosquitoes), and Formosus virus (from AWK and KIN mosquitoes). The C6/36 cells generated a strong viral piRNA response to all three viruses with remarkable resemblance to the primarily plus strand viral piRNA patterns observed in the cognate whole mosquito sample (**Fig. 8C**). The *Dcr2* mutations in C6/36 cells prevent these cells from generating conventional siRNAs from a double-stranded RNA (dsRNA) intermediate made during virus replication^41^, which could partially explain the lack of virus minus strand siRNAs. It is possible that apparent virus plus strand siRNAs in C6/36 cells could also be piRNAs that have been extensively trimmed to this shorter siRNA-like length.

The TMBTLV infection of Aag2 cells, which have an intact *Dcr2* gene, showed a clear pattern of viral siRNAs processed from a dsRNA intermediate since minus and plus strand siRNAs evenly covered the entire virus genome (**Fig. 8D**). This intact antiviral RNAi response in Aag2 cells against TMBTLV explains the slower infection kinetics in Aag2 cells versus C6/36 cells (**Fig. 7C**). However, Anphevirus and Formosus virus both did not generate antisense siRNAs in Aag2 cells, but instead produced mainly plus strand piRNAs from all the viral genes except the RdRP gene, in striking similarity to the small RNA patterns in the whole mosquito (**Fig. 8D**). Our results set the stage for utilizing these cell cultures and new mosquito virus stocks to study the biochemical mechanisms underlying the effective transmission of these viruses despite the triggering of small RNA responses in mosquito cells and animals.

## DISCUSSION

We present here the most comprehensive global mosquito virus small RNA survey to date and demonstrate how strikingly frequent are the robust small RNA responses to the persisting viruses within wild mosquitoes and in recently established colonies. In addition to analyzing new small and long RNA sequencing datasets, our MSRG pipeline also integrated previously published small RNA datasets^22^ to yield a more complete picture of the geographic distributions of mosquito viruses. Our analysis confirms the global reach of PCLV (and to a lesser extent HTV) infecting *Ae. aegypti*, but geographical boundaries of mosquito viruses based on small RNA profiles can still be observed such as virus delineations in the Americas continents and African-specific viruses like the Formosus virus. By comparing viral long RNAs to small RNAs, we uncovered new insights into the dynamics of virus replication intermediates that can evade RNAi interactions, mechanisms which we can explore in the future by infecting these viruses into mosquito cell cultures while knocking down mosquito host genes.

### Limitations and future opportunities of the study

Small RNA sequencing is a low throughput assay with more complicated library construction methods compared to long RNA sequencing. RNA sequencing is also less practical for routine virus surveillance conducted by municipal vector control organizations, which rely on primer panels in RT-PCR assays due to speed and low cost. Our current MSRG bioinformatics analysis is now more streamlined from the initial iteration^23,26^, but it remains a challenge to keep pace with the deluge of new virome entries and metagenomics studies^4,5,27^. Because of throughput limitations with small RNA sequencing scalability, our study is a snapshot in time and limited to specific locales despite our efforts to profile mosquitoes globally. Some of the narrow sampling issues are logistical and political restrictions on access to mosquitoes from certain countries.

It is possible that some of the viruses detected in colonized mosquitoes could result from a lab-acquired infection when multiple colonies share the same insectary space. Colonies that have been perpetuated for many generations may begin to reflect the environment that they are raised in instead of the locale that they originate from and are meant to represent. For instance, we detected the American-prevalent Anphevirus and CFAV in the Lambrechts lab’s Lope strain and the McBride lab’s ZIK strain, respectively. These African-originating strains had the most generations of lab propagation (>15 generations) compared to the other African colonies (average ∼8 generations, Fig. 8A).

Climate change is expanding the ecological niches for mosquitoes like *Ae. aegypti*, which have now invaded further into North America and Europe^35^. Broadening our small RNA genomics surveys to include animal captures in new locations would illuminate the potential disease vectoring threat that these invading mosquitoes would pose to human populations.

Small RNA sequencing provides extra rigor beyond the RT-PCR assay because the virus is expected to have replicated at high enough levels to provide the long RNA precursors needed for the mosquito RNAi machinery to convert into small RNAs. Thus, it was particularly surprising to observe the first case of DENV small RNAs in a wild mosquito isolate (**Fig. 4F**). It remains an open question whether wild mosquitoes displaying arboviral small RNAs are still actively infectious or if the RNAi response is now suppressing virus transmission. Only sequencing approaches, not RT-PCR, can discover new viruses and RNAi responses in mosquitoes, and our study demonstrates the utility of this approach to augment arbovirus surveillance programs.

### Could persistent insect viruses in wild mosquitoes be a source for emerging pathogens?

It is practical for academic lab mosquito strains to be clean of persistent viruses that would complicate genetic analyses. However, to better model arbovirus vector competence in the wild where human populations are affected, we need to study further the impact of persistent viruses that generate abundant viral small RNAs in mosquitoes. The infectivity of the mosquito viruses into cell cultures was surprising considering the apparent viral small RNA responses exhibited by the mosquito animals and the cell cultures. We did have occasional issues with mosquito viruses transferring between culture flasks by possible aerosolization within the biosafety cabinet during the handling of multiple mosquito homogenates (**Fig. 8**). This infectivity could explain the persistence and widespread detection of these viruses in so many of the mosquito samples profiled in this study.

An open question is whether persistent mosquito viruses can be transmitted from the mosquito to the bitten animal during blood feeding. For instance, it is unclear if massive vertical virus transmission of TMBTLV via maternal contribution to the eggs (**Fig. 6D**) would reflect a meaningful reservoir of viruses in the mosquito salivary glands and proboscis. On the other hand, the LTLV had very high levels of small RNAs in the midgut (**Fig. 3B, Supp. Fig. S4A**), suggesting this virus could likely mix with an initial bloodmeal and be transmitted during subsequent blood feedings. Notably, several of these mosquito viruses generating an RNAi response are related to plant viruses, suggesting that mosquito feeding on plant sap could be another modality of virus transmission, and perhaps these viruses are suppressing RNAi with factors related to P19 found in plant tombusviruses^34^. Lastly, our preliminary evidence shows that TMBTLV could be transmitted to mammalian Vero E6 and Huh7.5 cells. This surprising result was reproduced across multiple independent replications of this infection. This finding points to the possibility that arboviruses like DENV and ZIKV also started as mosquito-specific flaviviruses like CFAV until they gained the capability to infect humans to facilitate new avenues of viral transmission.

### Can the Formosus virus and other mosquito viruses affect arbovirus vector competence?

Previous studies on genetic variations causing lower ZIKV vector competence in *Ae. aegypti formosus* compared to *Ae. aegypti aegypti* were done prior to our new results on the Formosus virus persisting and being transmitted vertically in these African colonies^38^. It is possible that the *formosus-aegypti* crosses conducted in the previous study may have also transmitted Formosus virus, and insect virus interactions with arboviruses are documented. For example, co-infection with CFAV reduces DENV and ZIKV replication in cell culture and transmission in mosquitoes^46^. Our next priority will be to test how these mosquito viruses interact with DENV and ZIKV in co-infection experiments.

In addition, these mosquito viruses may also be useful agents for mosquito transduction and control in a similar vein as insect densoviruses^47,48^. Densoviruses have been engineered to carry reporter genes and toxic gene knockdown cassettes as prototype genetic tools to manipulate mosquitoes, but technical challenges must be overcome to realize the potential of densoviruses as mosquito genetic tools. We observed that, like the insect viruses investigated in this study, densoviruses also engage with the mosquito RNAi pathway to generate viral small RNAs (**Fig. 8B** and Ma, et al^23^). Rivaling densovirus in genome compactness is TMBTLV at just ∼4.5kb across two RNA segments. We are still characterizing the viral genomic RNAs’ 5’ and 3’ ends to build and test infectious clones, but there is potential for new mosquito genetic tools to be garnered from such a survey.

Previous studies have tried to harness RNAi to control mosquito populations and curtail arbovirus transmission^49–54^, but dsRNA delivery by injection has a low throughput, and RNAi by dsRNA ingestion is also limited in mosquitoes^55^. Mosquito viruses like TMBTLV that can readily trigger dsRNA and siRNAs in mosquitoes and cells (**Fig. 2B, 8D**) may solve the siRNA delivery problem. Future small RNA sequencing surveys like this can potentially discover new virus-vector biology, facilitating the development of new tools to combat the health threat that arboviruses pose to humans.

## MATERIALS AND METHODS

### Construction of small and total RNA libraries from whole mosquitoes and cells

Total RNA was extracted from whole frozen mosquitoes or mosquitoes preserved in RNA protection solutions. Zirconium beads (3.0mm) in a bead beater were used to homogenize mosquitoes before proceeding with NEB Monarch Total RNA Miniprep Kit (NEB #T2010). The on-column DNase I treatment was performed during RNA extraction. Small RNA libraries were made using NEBNext Small RNA Library Prep Set (NEB #E7330) with up to 5µg of RNA input. During library amplification, samples were put through up to 25 total PCR cycles. Total RNA libraries were made using Zymo-Seq RiboFree Total RNA Library Kit using up to 250ng RNA input, and following manufacturer’s protocol. All libraries were checked on an Agilent Bioanalyzer 2100 using either DNA 1000 or High Sensitivity DNA kit and sequenced at the BUMC Microarray and Sequencing Resource on an Illumina Next Seq 2000 using P3 flow cells for 50SE and 50PE reads for small RNA and total RNA libraries, respectively.

### Running VirusDetect with TPA and MAG database modifications

The source code for VirusDetect Version 1.7 ^28^ was downloaded from the GitHub repository (https://github.com/kentnf/VirusDetect). Additionally, the preprocessed VRL virus database Version 248 was obtained from the VirusDetect webpage (http://bioinfo.bti.cornell.edu/ftp/program/VirusDetect/virus_database/v248/). The source code and the corresponding database were deployed on the Shared Computing Cluster of Boston University. To meet VirusDetect requirements, the following packages were installed on the cluster: perl version 5.28.1, bioperl version 1.7.2, and python3 version 3.8.10.

The Third Party Annotation (TPA) GenBank files were sourced from the NCBI database (https://ftp.ncbi.nlm.nih.gov/tpa/tsa/). An in-house R pipeline was utilized to extract GenBank records, preserving 45,601 entries with the taxonomy ‘VIRUS’ for downstream processing. For each entry, both the genome sequence and corresponding protein sequences were retrieved from the NCBI database using the ACCESSION ID and saved in database files formatted for VirusDetect. Two id-mapping files were also generated following VirusDetect’s instructions. In addition to the GBVRL and TPA datasets, partial virus sequences in GenBank were annotated as Metagenome Assembled Genome (MAG). All MAG records downloaded from GenBank underwent the same processing steps as the TPA database. A total of 65,167 entries were retained to construct the modified VirusDetect database.

### Sequencing and bioinformatics analysis of small and long RNAs from mosquitoes

RNA Libraries were selected if they were above 1 million reads to run through the Mosquito Small RNA Genomics (MSRG) pipeline^23,26^. Outputs from MSRG include the alignment to a curated virus database. This includes breakdowns (in reads per million) of reads coming from siRNAs (18-23 nt) and piRNAs (24-35 nt), which strand reads map to, and a ratio of peaks to the average distance between peaks (used as a metric of coverage across viral sequence).

Some of the libraries in the Olmo et al study^22^ were made using a modified version of the NEBNext Multiplex Small RNA Library Prep Set where a random 6-mer was attached to the 5’ adapter. A Cutadapt trimming step in MSRG was adapted to remove the first six bases of each read. This modified pipeline was then used to run MSRG on these samples.

Total RNA libraries were sequenced as 50PE reads but were processed to look like small RNA reads so that they could also be run using the MSRG pipeline. All reads were trimmed to 35nt, R1 was reverse complemented to be in the same orientation as R2, and then the reverse complemented R1 was merged with R2. The resulting FASTQ file was then run through MSRG and processed just like the small RNA libraries.

Viruses were only considered to be present in a sample if it had at least 10 reads per million (RPM) and a coverage ratio greater than 0.75. Additionally, a strand bias score was calculated for each virus/sample pair by taking the ratio of reads mapping to the top versus bottom strand. For plots made using R/RStudio, packages used include: ggplot2, tidyverse, and ggpubr.

In the bubble plot, log10 of the strand bias score was mapped to the color, and log10 of the RPM was mapped to the size. For the strand bias, any value greater than 1 (a greater than 10-fold strand bias) was squished to be the “max” color on the scale. For the scatterplots comparing the small and total RNAs, the 10 RPM ratio was ignored in order to completely compare the two libraries. Sample/virus pairs were kept if at least one of the two libraries was at least 10 RPM.

### Cell Cultures

C6/36 and Aag2-TC mosquito cell cultures were propagated in DMEM (Gibco) supplemented with 10% FBS and 1% Tryptose Phosphate Broth (TPB) and maintained at 28° C and 5% CO2. HUH7.5 and Vero E6 TC cell cultures were propagated in DMEM (Gibco) supplemented with 10% FBS and 1% P/S (Penicillin/Streptomycin) and maintained at 37° C and 5% CO2. The mammalian cell cultures were split by washing with 1X PBS and 0.25% Trypsin EDTA.

### Cell infection with mosquito viruses

The mosquitoes received from Awka (Nigeria), Kintampo (Ghana), Pozo Rica (Mexico), and Banzon Research (USA) were used to infect the cell lines with viruses. Mosquitoes were homogenized in 1.5 mL DMEM with zirconium beads (3.0mm) in a bead beater. The homogenate was passed through a 0.45µm pore size filter and added into a T-25 flask containing ∼90% cell confluency in 5ml of media in T-25 flasks. On the first day of infection only, a cocktail of Primocin, Normocin, and Fungin antibiotics (InvivoGen) was added to the infection media.

For mosquito cell blind passage, the medium was collected from the infected cells, passed through the 0.45µm pore size filter, and added into a new T-25 flask containing ∼90% cell confluency. For serial passage, the cells infected by homogenizing the mosquitoes were split into a new flask with a fresh medium. This process was repeated every 7th day for both blind and serial passage. The cells collected on the 7th day were used for total RNA extraction to confirm the virus infection.

For mammalian cell blind passage experiments, VeroE6 and Huh7.5 cells were infected with non-titered virus stock medium collected from C6/36 cells infected with the mosquito-viruses from Awka, Nigeria (AWK) and Benzon Research (BZL). The collected medium was passed through a 0.45µm pore size filter and was added to a new T25 flask containing ∼90% confluency. On the 4th day post-infection, the medium was replaced with a fresh batch of virus stock medium. On the 7th day, medium from the cells was transferred out and cells were washed twice with 1X PBS before trypsinizing cells for total RNA extraction and archiving live cells by cryopreservation. This process was repeated on every 7th day.

### RNA extraction and RT-qPCR analysis of virus amplicons

Total RNA was extracted with Monarch Total RNA Miniprep Kit (NEB) following the manufacturer’s instructions, including the DNase I treatment step. For RT-PCR, 1µg RNA was used for first strand cDNA preparation by using ProtoScript II Reverse Transcriptase kit followed by Phusion High-Fidelity DNA Polymerase (NEB). RT-qPCR was performed with LunaScript RT Supermix (NEB) in a Bio-Rad CFX Opus 96 Real-Time PCR System. The housekeeping genes, *Ae. aegypti* actin (GenBank accession XM_001659913) and *Ae. albopictus* actin (GenBank accession XM_019702203), were used as reference genes to normalize the target gene expression by 2-ΔΔCt methods.

### Virus Kinetics

The virus kinetics of TMBTLV, Formosus virus, and Totivirus were measured in C6/36 and Aag2-TC cell lines. First, the virus particles were quantified in the virus stock medium by digital-droplet PCR on the Bio-Rad QX-200 instrument. The cells were infected with 5 mL virus stock medium in the T-25 flask. The cells were collected from the flask on the 1st, 3rd, 5th, 7th, and 10th day post-infection. At each time point, the cells were resuspended in the medium using a pipette, and one mL of the medium containing the cells was replaced with a fresh medium. The cells were collected and processed from one mL for the total RNA extraction. Subsequently, the virus quantification was done by RT-qPCR, as described above. This experiment was repeated in three independent biological replicates for each time point.

### Resource availability Lead contact

Further information and requests for resources and reagents should be directed to and will be fulfilled by the lead contact, Nelson Lau.

### Materials availability

All unique/stable reagents generated in this study are available from the lead contact. Material transfer agreements with Boston University may apply.

## Data and code availability

All sequencing data produced and generated by this study is available on Sequencing Read Archive (SRA) under BioProject PRJNA1104658. See Tables S1 and S2 for specific BioSample and SRA accessions. SRA accessions for publicly available datasets used in this study can be found in Table S1C. The MSRG pipeline code can be found on the Github repository: https://github.com/laulabbumc/MosquitoSmallRNA.

## ACKNOWLEDGEMENTS

We thank Mohsan Saeed and Fabiana Feitosa-Suntheimer for comments on this manuscript. We acknowledge Anubis Vega-Rúa and Silvânia da Veiga Leal as the source of mosquito colonies from Guadeloupe and Cape Verde, respectively. We thank João Marques for assistance in accessing his lab’s public datasets. We acknowledge Mark Stenglein and Marylee Kapuscinski for technical assistance in small RNA sequencing.

N.C.L.’s lab was funded by NIH/NIGMS (GM135215). L.L. was supported by the French Government’s Investissement d’Avenir program, Laboratoire d’Excellence Integrative Biology of Emerging Infectious Diseases (grant ANR-10-LABX-62-IBEID). A.B. was supported by a stipend from the Pasteur - Paris University (PPU) International PhD Program. A.E.W. and K.E.O. were supported by R01 AI130085. A.E.W. and E.C. research was supported by the Intramural Research Program of NIH/NIAID (AI001246). N.H.R and C.S.M.’s work here was supported by NIH GRANT R00DC012069 and a New York Stem Cell Foundation Robertson Neuroscience Investigator Award. A.L.C.S. and M.D were supported by U.S. Centers for Disease Control and Prevention (CDC) Cooperative Agreement Number 1U01CK000510, Southeastern Regional Center of Excellence in Vector-Borne Diseases Gateway Program. The CDC did not have a role in the design of the study or the collection, analysis, or interpretation of data. D.E.B.’s work was supported in part by grants from the National Institutes of Health, National Institute of Allergy and Infectious Diseases (AI148477). M.A.Y. is supported by the Searle Scholars Program, the Richard and Susan Smith Family Foundation, the Esther A. & Joseph Klingenstein Fund, and the Simons Foundation.

## Author contributions

Conceptualization, N.C.L, S.G., R.S., A.E.W., and Z.Z.; Methodology/Investigation, N.C.L, S.G., R.S., A.C.V., A.E.W., Z.Z., and G.D.; Formal Analysis, S.G., N.C.L., and R.S.; Data Curation/Software, C.Z., S.G., and Z.Z.; Writing – Original Draft, N.C.L., S.G., and R.S.; Writing – Review & Editing, N.C.L., S.G., A.E.W., L.L., N.H.R., C.S.M., and Z.Z.; Visualization, N.C.L., S.G., R.S., and Z.Z.; Funding Acquisition, N.C.L., L.L., K.E.O., E.C., C.S.M., M.A.Y., D.E.B., and M.D.; Sample Contribution & Resources, A.E.W, I.S.V., N.H.R., A.G.S., D.E.B., J.M., S.S.W., A.C., T.R., M.S., A.B., J.A., O.B.A., D.A., W.L.L., C.H.C., C.V., C.G.A., A.P., T.M., B.C., D.W., D.S., M.A.Y., A.L.C.S., M.D., A.B., L.L., C.S.M, K.E.O., and E.C.

## FIGURES and TABLES LEGENDS

**Supplemental Figure S1. Using the AEFE1 EVE as a constant quality tracker control for viral small RNA profiling and implementing VirusDetect to discover new ISVs from small RNAs.**

(A) Coverage plots on left and bubble plots on right representing the small RNAs from the Endogenous Viral Element (EVE) from *Ae. aegypti* called AEFE1 (Suzuki et al, 2017). (B) The results output table from the VirusDetect program only loaded with the GBVRL database while analyzing the small RNA library from one of the BZL lab strain samples. (C) Coverage of the de-novo small RNA assembly contigs generated by VirusDetect of the corresponding virus determinations from the table in (B). The red asterisks mark a false-positive call against a tombus-like virus entry in GBVRL with limited contig coverage. (D) VirusDetect results analyzing the same BZL lab strain small RNA library in (B) but now with a custom loading of the TPA and MAG databases that are distinct from GBVRL. This result now shows two new viruses with much more complete contig coverage in (E), such as the Tiger Mosquito Bi-segmented Tombus-Like virus (TMBTLV) that is the true hit replacing the false-positive call in (B-C).

**Supplemental Figure S2. Read length distribution profiles of all the *Ae. aegypti* small RNA libraries analyzed in this global survey.**

Read length distribution plots as percentages of the libraries with functional classes of reads represented by different colored lines. The groups correspond to (A) Lab strains, (B) North American strains, (C) Central and South American strains, (D) Asian strains, (E) African strains, (F) Cell lines, (G) Analyzed-separately-as-additional-replicates samples.

**Supplemental Figure S3. Complete bubble plots of viral small RNAs sampled in this global survey study.**

Top plots are the siRNA-length small RNAs (18-23nt), the middle plots are the piRNA-length small RNAs (24-35nt), and the bottom plots are all the small RNAs as shown in the main figures 2 – 6. The groups correspond to (A) Lab strains, (B) North American strains, (C) Central and South American strains, (D) Asian strains, and (E) African strains. (F) are additional samples that we considered as replicates of other libraries but did not include in the main study because of concerns of potential cross-sample contamination.

**Figure S4. Plant-related viruses and other prevalent viruses in American *Aedes aegypti* shared on both east and west coasts.**

(A-B) Coverage plots of two insect viruses related to plant-originating viruses marked in Figure 3. (C) Maps marking approximate locations of *A. aegypti* samples collected from California, Louisiana and Florida. Coverage plots of viral small RNAs from whole *A. aegypti* from: (D) *Aedes* Binegev-like virus-Cayenne strain from California and Florida; and (E) Renna virus from three states. (F) Principal Component Analysis of virus small RNA levels can discriminate different groups of mosquito samples in between the North American, Central & South American, and Lab strains.

**Figure S5. Tracking virus presence with RT-PCR during blind passaging of C6/36 mosquito cells from infections with PZR, AWK and KIN mosquito homogenates – with the potential to infect mammalian cells.**

(A) RT-PCR of viral amplicons against the Verdadero virus RdRP and capsid genes. Two independent infections with two batches of PZR mosquitoes. (B) RT-PCR of viral amplicons against Tesano virus, Totivirus and Formosus virus from two independent infections with two batches of AWK and KIN mosquitoes. The actin control are primers against *Ae. albopictus* actin. (C) Continued blind passaging of virus stocks from C6/36 cells are then tested for infection of mammalian VeroE6 and Huh7.5 cells followed by RT-PCR detection of viral amplicons. Red arrowheads point to insect virus amplicons detected in the mammalian cells. Asterisk marks a totivirus already present in this lab stock of C636 cells. (D) Brightfield images cells from after 1 week of infection from 4^th^ passage in (C) before harvesting total RNA for RT-PCR analysis. The cytopathic effect of infection from the BZL virus stock is most evident on C6/36 and Huh7.5 cells.

**Figure S6. Cloning and sequencing the TMBTLV and Formosus virus from *Ae. aegypti*.**

(A) Contig assembly of Sanger sequencing of cloned TMBTLV amplicons from S1 and S2 RNAs. (B) Diagrams of SNP mutations from our cloned fragment sequencing versus the initial TMBTLV references BK059489 and BK059490. (C) Contig assembly of Nanopore sequencing of cloned Formosus virus amplicons. (D) Diagrams of SNP mutations from our cloned fragment sequencing versus the initial Formosus references BK059424. Both of our TMBTLV and Formosus virus variants sequences have now been contributed to GenBank with verified sequence accessions that should facilitate its inclusion in the future updates of the GBVRL database.

**Table S1. Overview of small RNA libraries made and used for this study.**

**Tab S1a. Mosquitoes. Tab S1b. Cell culture infections. Tab S1c. Previously published datasets.**

**Table S2. Overview of total RNA libraries made and used for this study.**

**Tab S2a. Mosquitoes. Tab S2b. Cell culture infections.**

**Table S3. Current List of Insect viruses in the MSRG pipeline analyzed in this study.** At the bottom are also some additional viruses that were circumspect, were either already removed or are being considered for removal in future extensions of this study.

**Table S4. List of oligonucleotides used in this study.**

