## Supplemental Figure S1 for "Global genomics of *Aedes aegypti* unveils widespread and novel infectious viruses capable of triggering a small RNA response"

Gupta, Sharma, et al.
**Figure S1.**

**A**

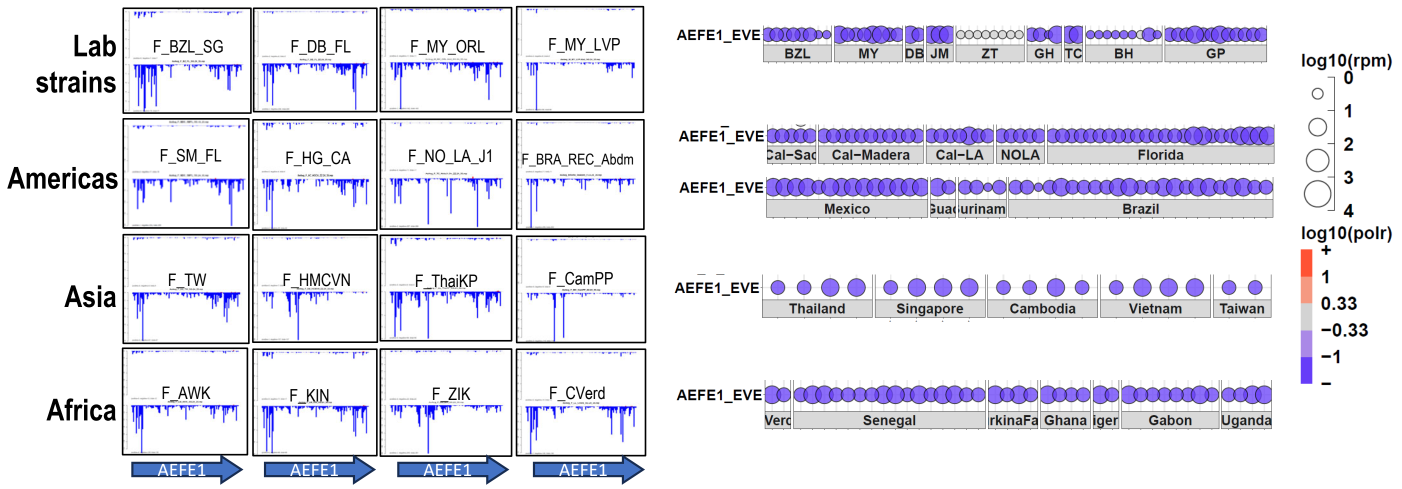


**B**

### VirusDetect result from Genbank Virus Reference List (GBVRL)

| Reference | Length | Coverage (%) | #contig | Depth | Depth (Norm) | %Identity | %Iden Max | %Iden Min | Genus | Description |
|---|---|---|---|---|---|---|---|---|---|---|
| MN053721 | 7582 | 5033 (66.4) | 54 | 27.3 | 0.4 | 98.65 | 100 | 95.00 | NA | Aedes aegypti toti-like virus strain 2017-Ab-AAF-1-5 capsid (orf1) and RNA-dependent RNA polymerase (orf2) genes, complete cds. |
| MH324433 * | 1925 | 429 (22.3) | 6 | 1066.8 | 17.4 | 89.30 | 97.92 | 83.15 | NA | Hypsignathus monstrosus tombus-like virus 1 isolate HMTLV1-RML51 hypothetical protein 1 and hypothetical protein 2 genes, complete cds. |
| KR003802 | 1609 | 1372 (85.3) | 7 | 8.8 | 0.1 | 98.69 | 100 | 97.70 | NA | Humaita-Tubiacanga virus isolate Rio segment capsid Capsid gene, complete cds. |
| KR003785 | 1358 | 547 (40.3) | 7 | 291.5 | 4.7 | 95.64 | 99.16 | 92.52 | phasivirus | Phasi Charoen-like virus isolate Rio segment S Nucleocapsid gene, complete cds. |
| MZ560693 | 148 | 143 (96.6) | 1 | 898.6 | 14.6 | 100 | 100 | 100 | picobirnavirus | Picobirnavirus sp. isolate PBV/Cryptosporidium/NZL/NZ03_CR_S1/2019 RNA-dependent RNA polymerase (RdRp) gene, partial cds. |
| MT815904 | 391 | 194 (49.6) | 3 | 733.6 | 11.9 | 92.31 | 93.75 | 91.23 | pospiviroid | Citrus exocortis viroid isolate CEYVd OH19-2, partial sequence. |
| MN225577 | 927 | 223 (24.1) | 1 | 202.5 | 3.3 | 97.31 | 97.31 | 97.31 | vesiculovirus | UNVERIFIED: Guampa vesiculovirus RNA-dependent RNA polymerase protein-like gene, partial sequence. |

**C**

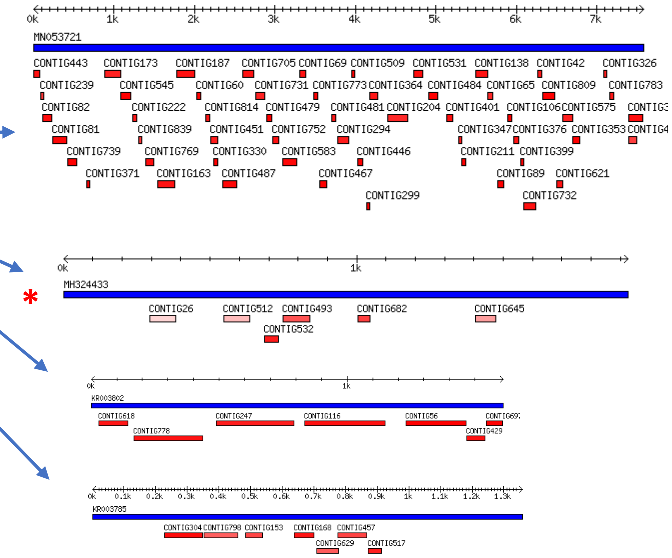


**D**

### VirusDetect result from TPA and MAG databases

| Reference | Length | Coverage (%) | #contig | Depth | Depth (Norm) | %Identity | %Iden Max | %Iden Min | Genus | Description |
|---|---|---|---|---|---|---|---|---|---|---|
| BK059430 | 1494 | 1270 (85) | 12 | 23.8 | 0.4 | 98.58 | 100 | 97.10 | TPA_asm: Aedes partiti-like virus 1 isolate Jane RNA-dependent RNA polymerase (RdRP) gene, complete cds | 1 |
| BK059431 | 1313 | 1275 (97.1) | 2 | 129.5 | 2.1 | 98.21 | 100 | 98.02 | TPA_asm: Aedes partiti-like virus 1 isolate Jane capsid protein gene, complete cds | 1 |
| BK059489 | 2338 | 2306 (98.6) | 2 | 2774.2 | 45.2 | 98.40 | 98.49 | 97.61 | TPA_asm: Tiger mosquito bi-segmented tombus-like virus isolate Phu Hoa HSP90-containing protein and RdRP genes, complete cds | 1 |
| BK059490 | 1903 | 1897 (99.7) | 1 | 5922.4 | 96.5 | 98.68 | 98.68 | 98.68 | TPA_asm: Tiger mosquito bi-segmented tombus-like virus isolate Phu Hoa hypothetical protein and putative structural protein genes, complete cds | 1 |

**E**

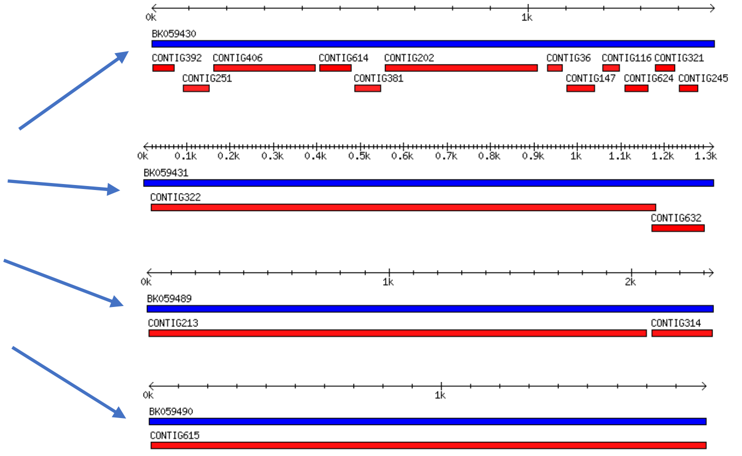


**Supplemental Figure S1. Using the AEFE1 EVE as a constant quality tracker control for viral small RNA profiling and implementing VirusDetect to discover new ISVs from small RNAs.** (A) Coverage plots on left and bubble plots on right representing the small RNAs from the Endogenous Viral Element (EVE) from *Ae. aegypti* called AEFE1 (Suzuki et al, 2017). (B) The results output table from the VirusDetect program only loaded with the GBVRL database while analyzing the small RNA library from one of the BZL lab strain samples. (C) Coverage of the de-novo small RNA assembly contigs generated by VirusDetect of the corresponding virus determinations from the table in (B). The red asterisks mark a false-positive call against a tombus-like virus entry in GBVRL with limited contig coverage. (D) VirusDetect results analyzing the same BZL lab strain small RNA library in (B) but now with a custom loading of the TPA and MAG databases that are distinct from GBVRL. This result now shows two new viruses with much more complete contig coverage in (E), such as the Tiger Mosquito Bi-segmented Tombus-Like virus (TMBTLV) that is the true hit replacing the false-positive call in (B-C).
