## Supplemental Figure S2 for "Global genomics of *Aedes aegypti* unveils widespread and novel infectious viruses capable of triggering a small RNA response"

**Gupta, Sharma, et al.**
**Figure S2.**

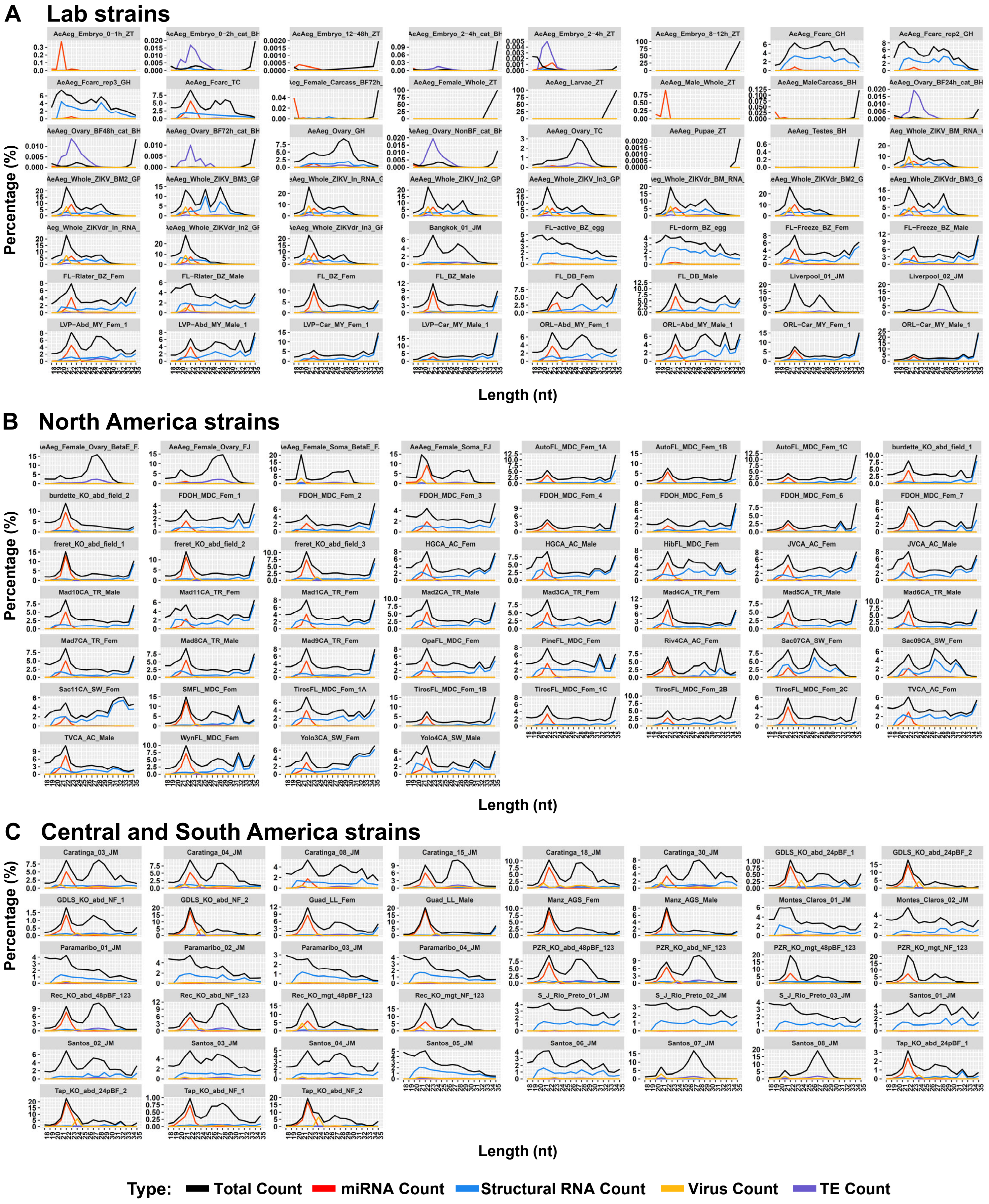


**Supplemental Figure S2. Read length distribution profiles of all the *Ae. aegypti* small RNA libraries analyzed in this global survey.** Read length distribution plots as percentages of the libraries with functional classes of reads represented by different colored lines. The groups correspond to (A) Lab strains, (B) North American strains, (C) Central and South American strains, (D) Asian strains, (E) African strains, (F) Cell lines, (G) Analyzed-separately-as-additional-replicates samples.

Gupta, Sharma, et al.
Figure S2.

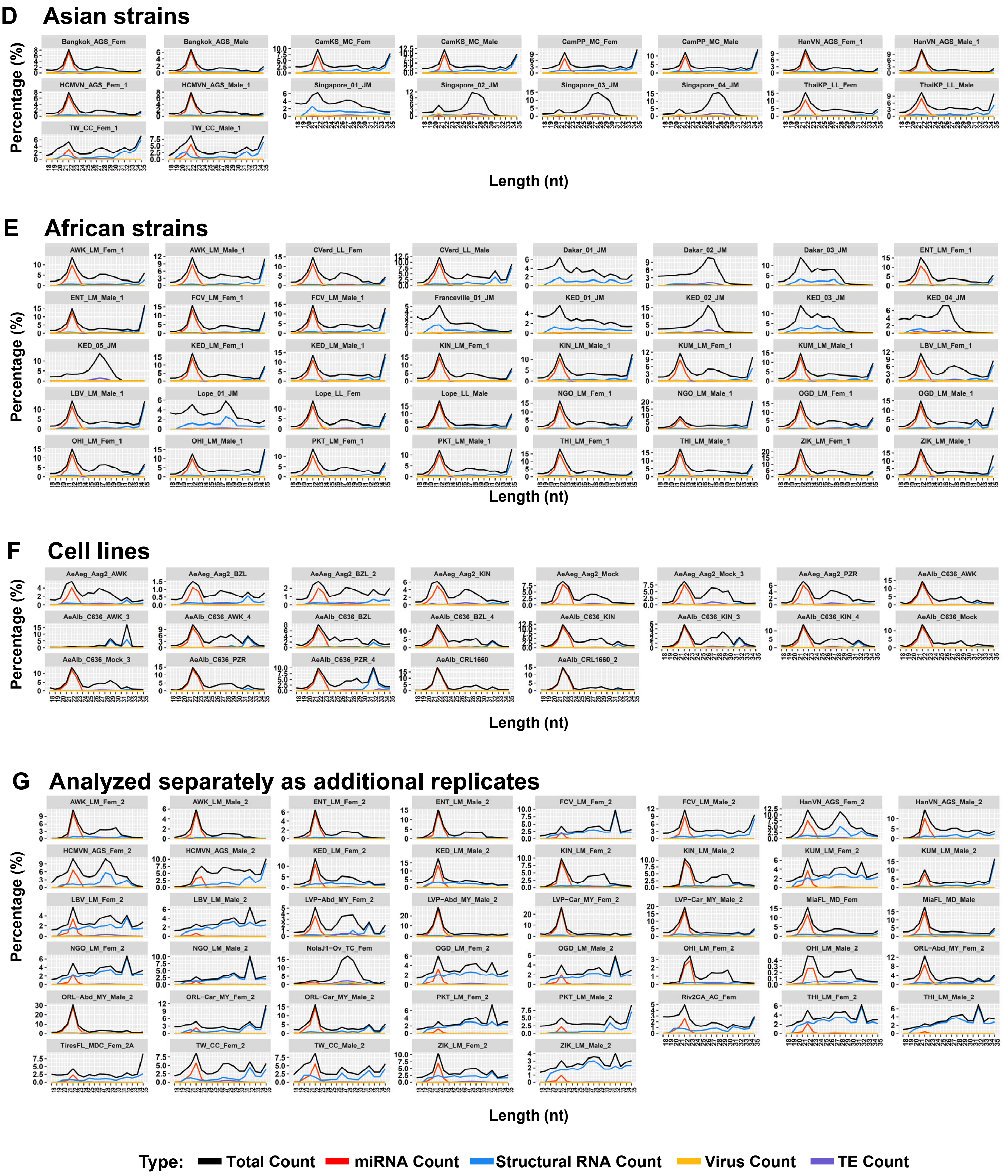


**Supplemental Figure S2. Read length distribution profiles of all the *Ae. aegypti* small RNA libraries analyzed in this global survey.** Read length distribution plots as percentages of the libraries with functional classes of reads represented by different colored lines. The groups correspond to (A) Lab strains, (B) North American strains, (C) Central and South American strains, (D) Asian strains, (E) African strains, (F) Cell lines, (G) Analyzed-separately-as-additional-replicates samples.
