## Supplemental Figure S3 for "Global genomics of *Aedes aegypti* unveils widespread and novel infectious viruses capable of triggering a small RNA response"

**Gupta, Sharma, et al.**
**Figure S3.**

**A**

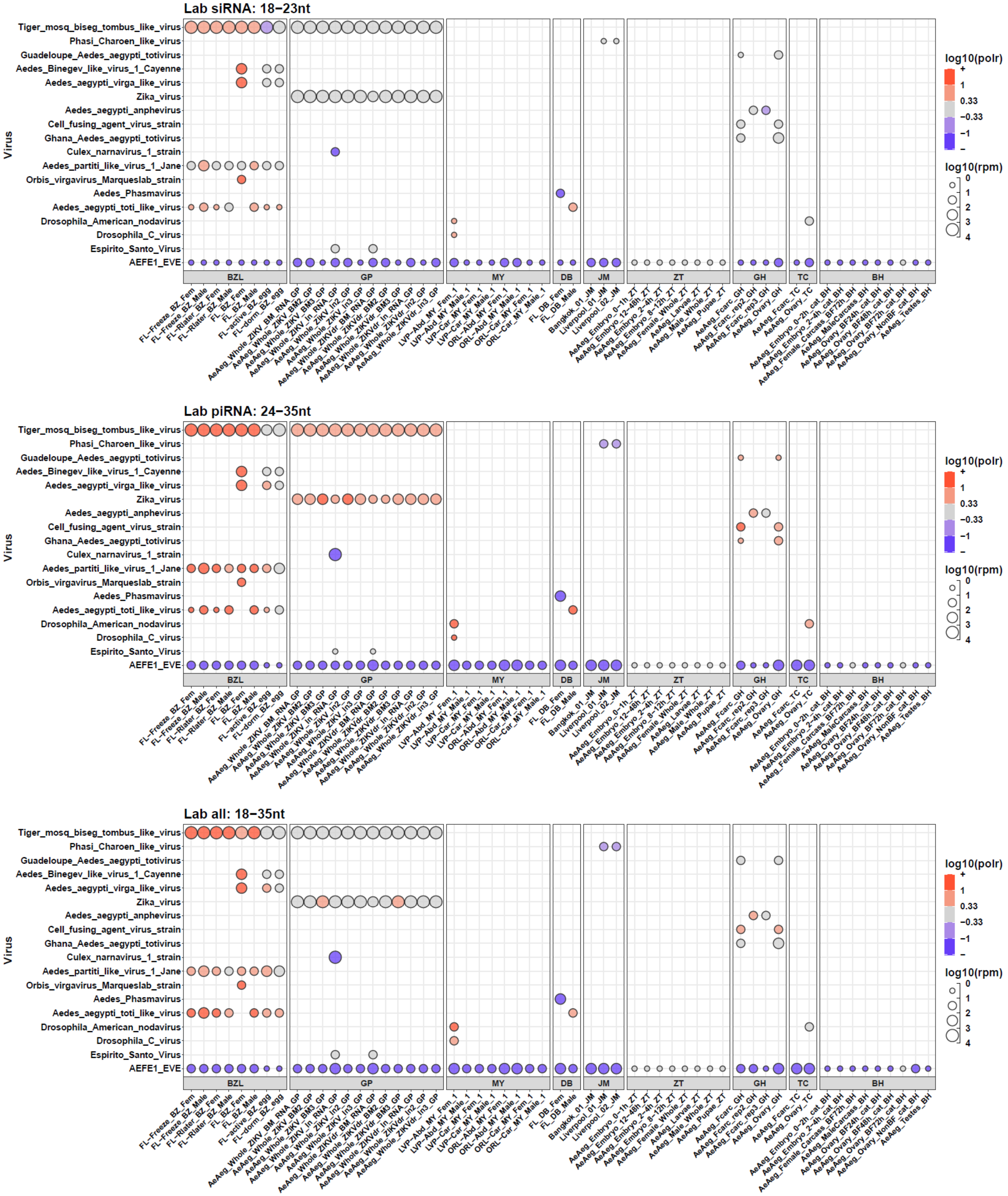


**Supplemental Figure S3. Complete bubble plots of viral small RNAs sampled in this global survey study.**
Top plots are the siRNA-length small RNAs (18-23nt), the middle plots are the piRNA-length small RNAs (24-35nt), and the bottom plots are all the small RNAs as shown in the main figures 2 – 6. The groups correspond to (A) Lab strains, (B) North American strains, (C) Central and South American strains, (D) Asian strains, and (E) African strains. (F) are additional samples that we considered as replicates of other libraries but did not include in the main study because of concerns of potential cross-sample contamination.

**Gupta, Sharma, et al.**
**Figure S3.**

**B**

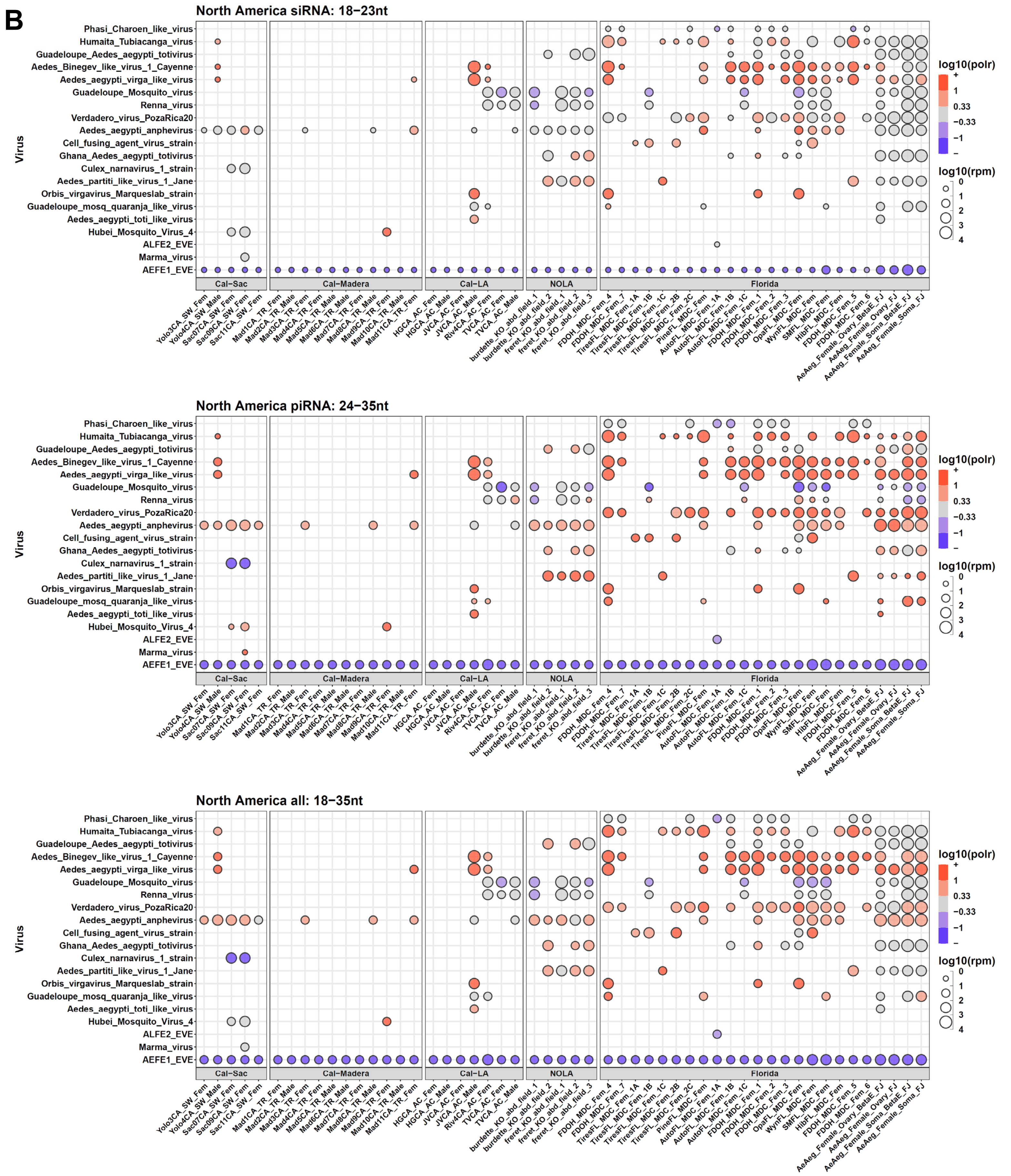


**Supplemental Figure S3. Complete bubble plots of viral small RNAs sampled in this global survey study.**
Top plots are the siRNA-length small RNAs (18-23nt), the middle plots are the piRNA-length small RNAs (24-35nt), and the bottom plots are all the small RNAs as shown in the main figures 2 – 6. The groups correspond to (A) Lab strains, (B) North American strains, (C) Central and South American strains, (D) Asian strains, and (E) African strains. (F) are additional samples that we considered as replicates of other libraries but did not include in the main study because of concerns of potential cross-sample contamination.

Gupta, Sharma, et al.
Figure S3.

C

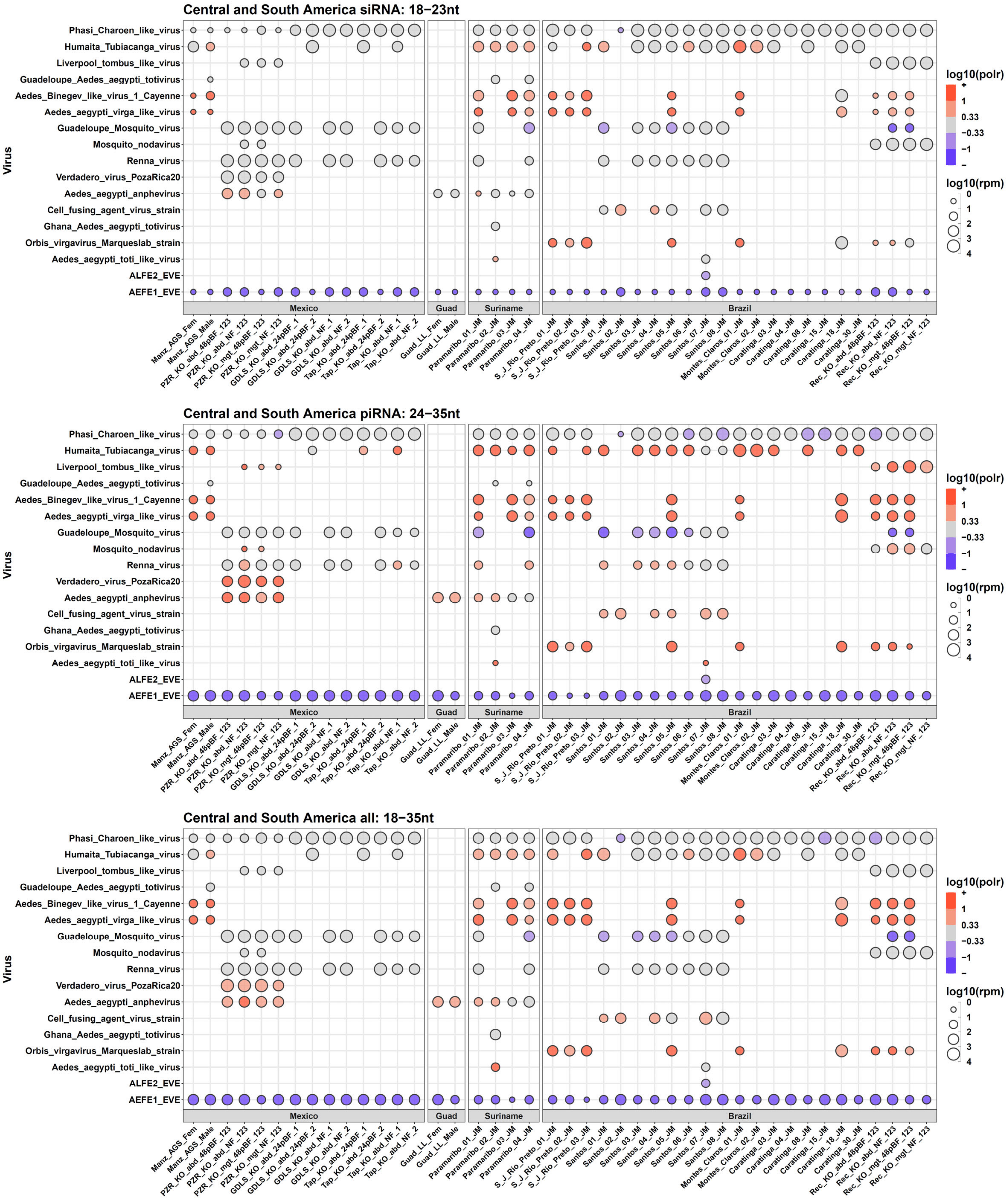


**Supplemental Figure S3. Complete bubble plots of viral small RNAs sampled in this global survey study.** Top plots are the siRNA-length small RNAs (18-23nt), the middle plots are the piRNA-length small RNAs (24-35nt), and the bottom plots are all the small RNAs as shown in the main figures 2 – 6. The groups correspond to (A) Lab strains, (B) North American strains, (C) Central and South American strains, (D) Asian strains, and (E) African strains. (F) are additional samples that we considered as replicates of other libraries but did not include in the main study because of concerns of potential cross-sample contamination.

**Gupta, Sharma, et al.**
**Figure S3.**

**D**

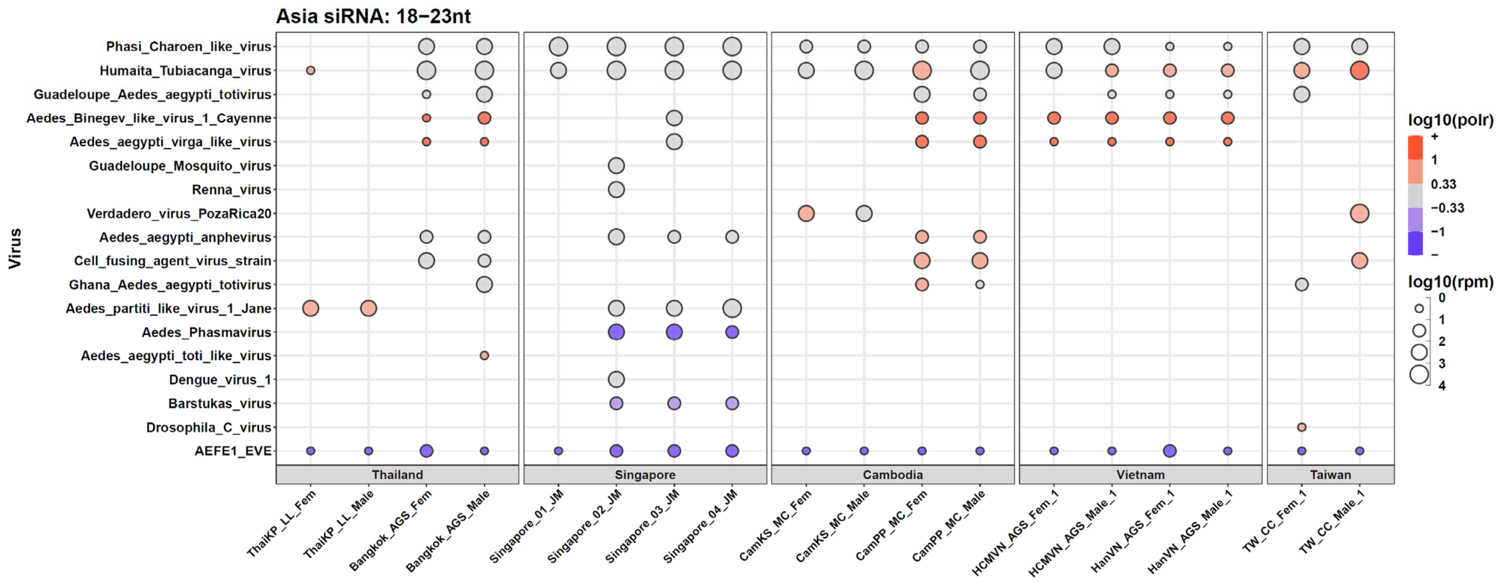


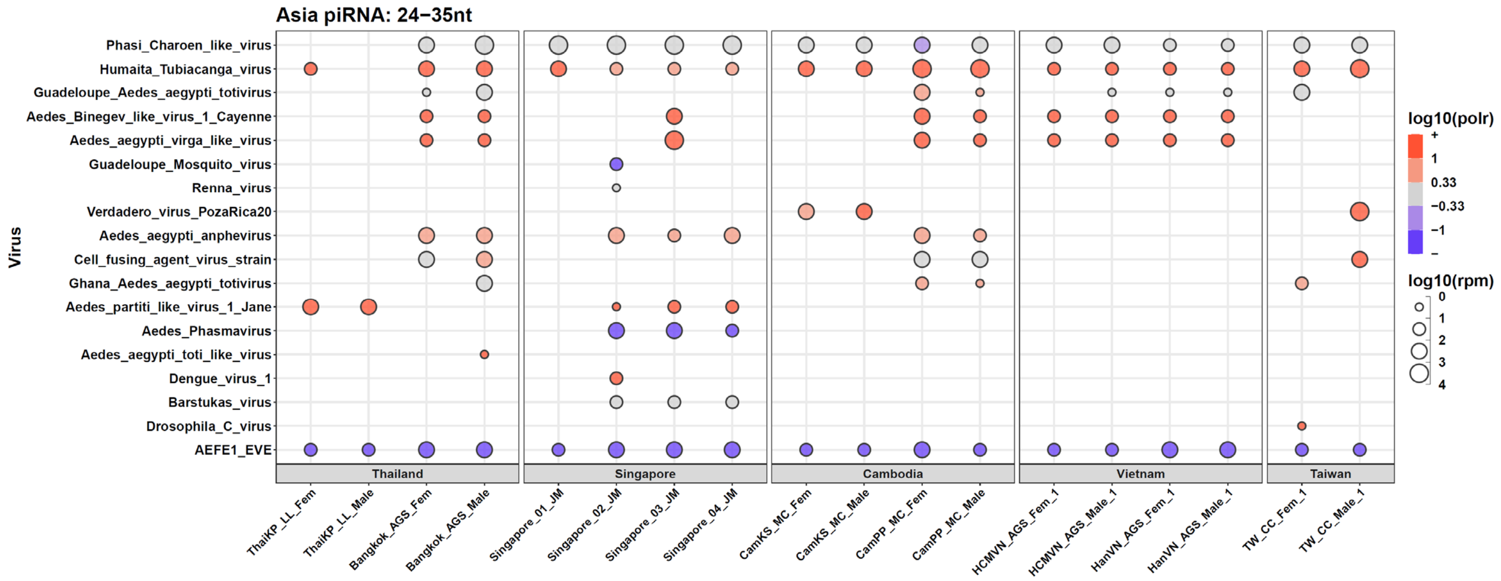


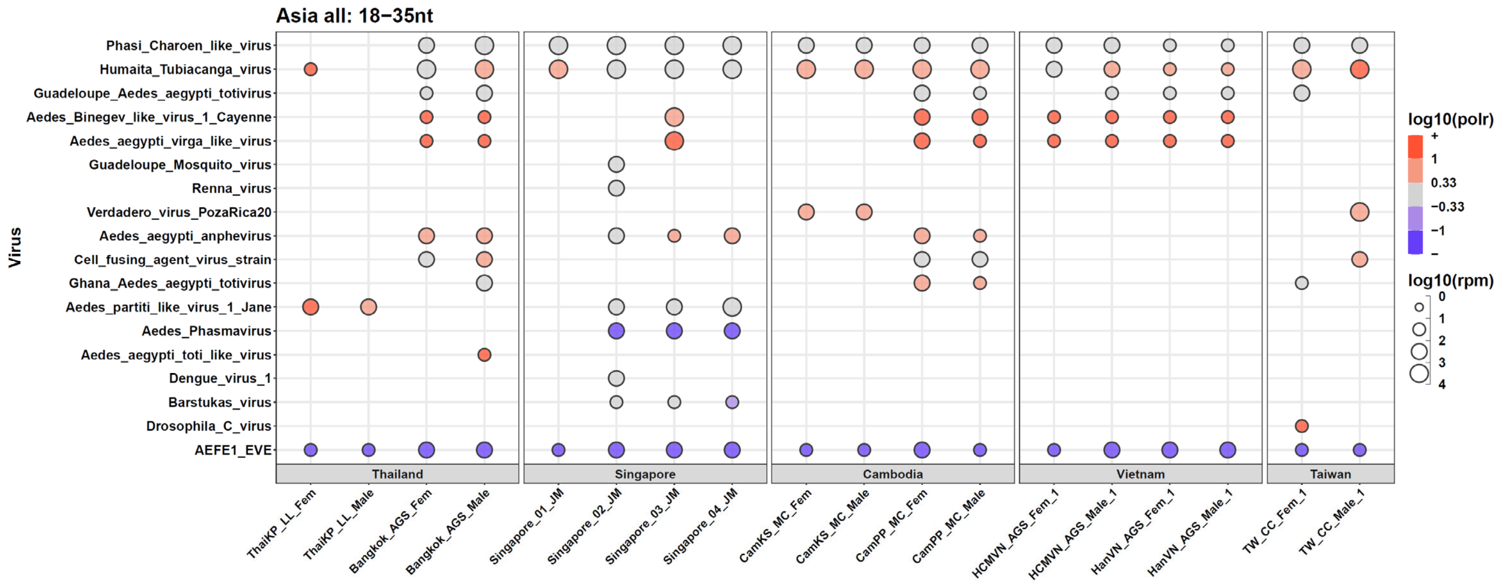


**Supplemental Figure S3. Complete bubble plots of viral small RNAs sampled in this global survey study.**
Top plots are the siRNA-length small RNAs (18-23nt), the middle plots are the piRNA-length small RNAs (24-35nt), and the bottom plots are all the small RNAs as shown in the main figures 2 – 6. The groups correspond to (A) Lab strains, (B) North American strains, (C) Central and South American strains, (D) Asian strains, and (E) African strains. (F) are additional samples that we considered as replicates of other libraries but did not include in the main study because of concerns of potential cross-sample contamination.

**Gupta, Sharma, et al.**
**Figure S3.**

**E**

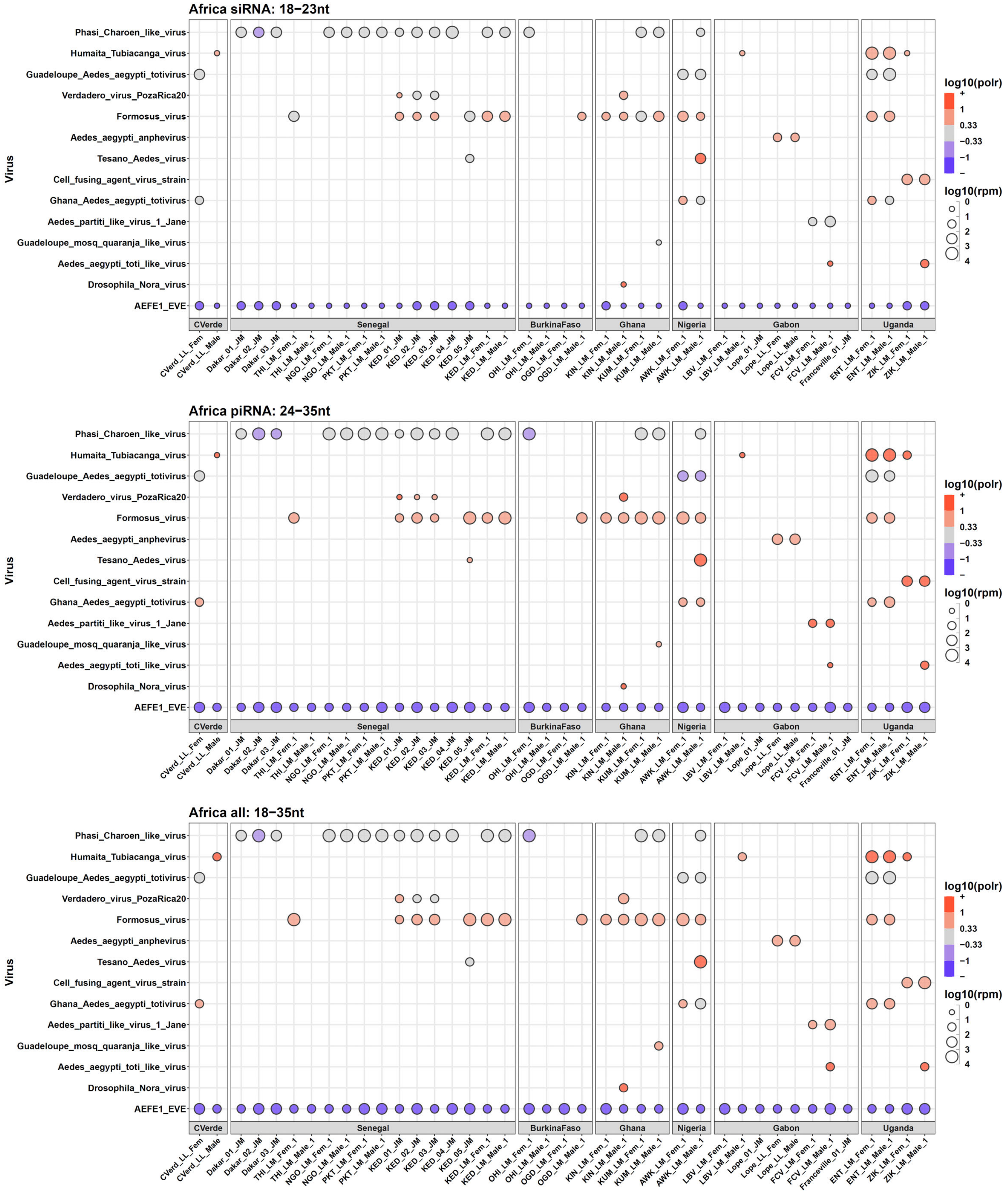


**Supplemental Figure S3. Complete bubble plots of viral small RNAs sampled in this global survey study.**
Top plots are the siRNA-length small RNAs (18-23nt), the middle plots are the piRNA-length small RNAs (24-35nt), and the bottom plots are all the small RNAs as shown in the main figures 2 – 6. The groups correspond to (A) Lab strains, (B) North American strains, (C) Central and South American strains, (D) Asian strains, and (E) African strains. (F) are additional samples that we considered as replicates of other libraries but did not include in the main study because of concerns of potential cross-sample contamination.

**Gupta, Sharma, et al.**
**Figure S3.**

**F**

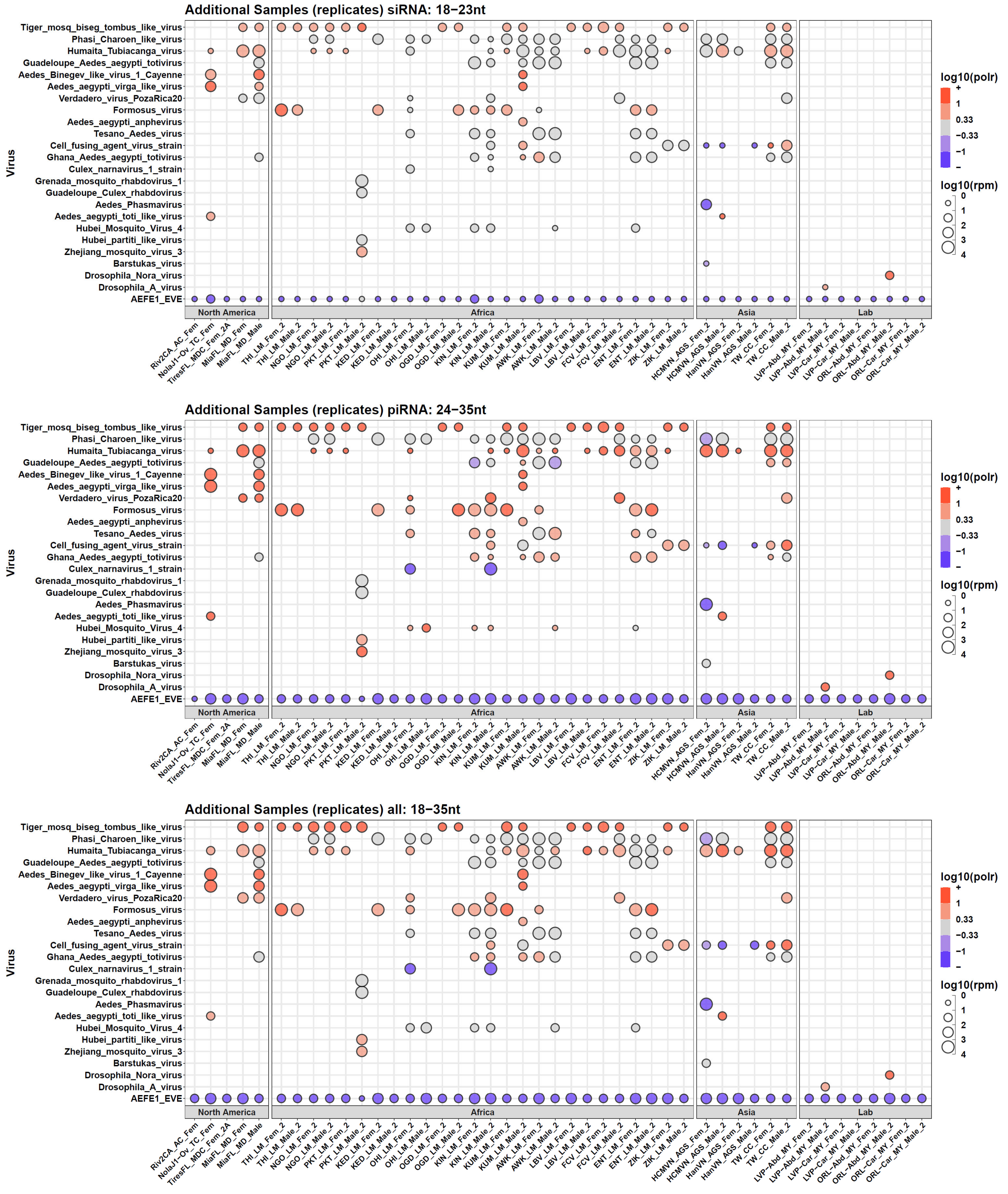


**Supplemental Figure S3. Complete bubble plots of viral small RNAs sampled in this global survey study.**
Top plots are the siRNA-length small RNAs (18-23nt), the middle plots are the piRNA-length small RNAs (24-35nt), and the bottom plots are all the small RNAs as shown in the main figures 2 – 6. The groups correspond to (A) Lab strains, (B) North American strains, (C) Central and South American strains, (D) Asian strains, and (E) African strains. (F) are additional samples that we considered as replicates of other libraries but did not include in the main study because of concerns of potential cross-sample contamination.
