## Supplemental Figure S4 for "Global genomics of *Aedes aegypti* unveils widespread and novel infectious viruses capable of triggering a small RNA response"

**Gupta, Sharma, et al.**
**Figure S4.**

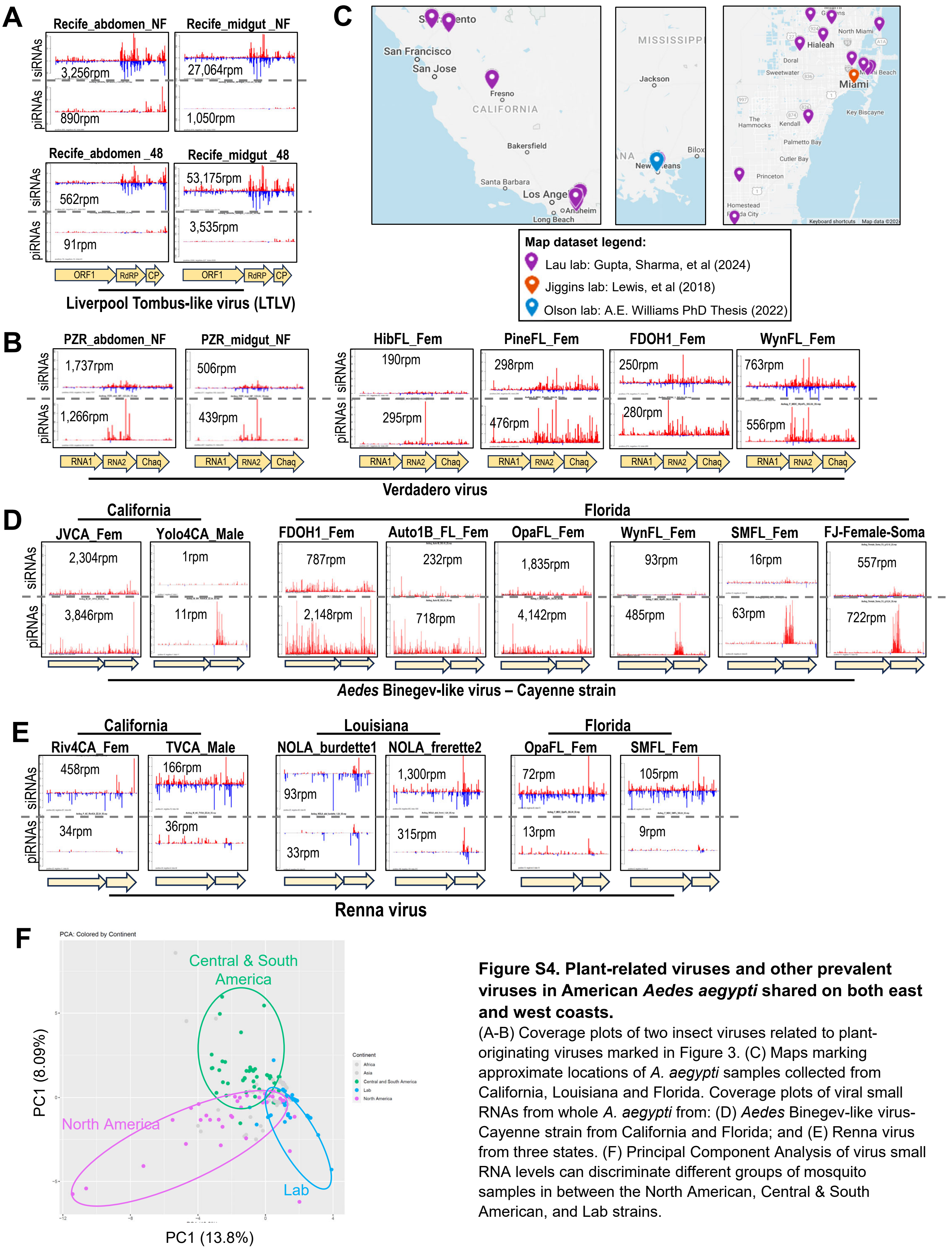


**Figure S4. Plant-related viruses and other prevalent viruses in American *Aedes aegypti* shared on both east and west coasts.**
(A-B) Coverage plots of two insect viruses related to plant-originating viruses marked in Figure 3. (C) Maps marking approximate locations of *A. aegypti* samples collected from California, Louisiana and Florida. Coverage plots of viral small RNAs from whole *A. aegypti* from: (D) *Aedes* Binegev-like virus-Cayenne strain from California and Florida; and (E) Renna virus from three states. (F) Principal Component Analysis of virus small RNA levels can discriminate different groups of mosquito samples in between the North American, Central & South American, and Lab strains.
