## Supplemental Figure S5 for "Global genomics of *Aedes aegypti* unveils widespread and novel infectious viruses capable of triggering a small RNA response"

Gupta, Sharma, et al.
Figure S5.

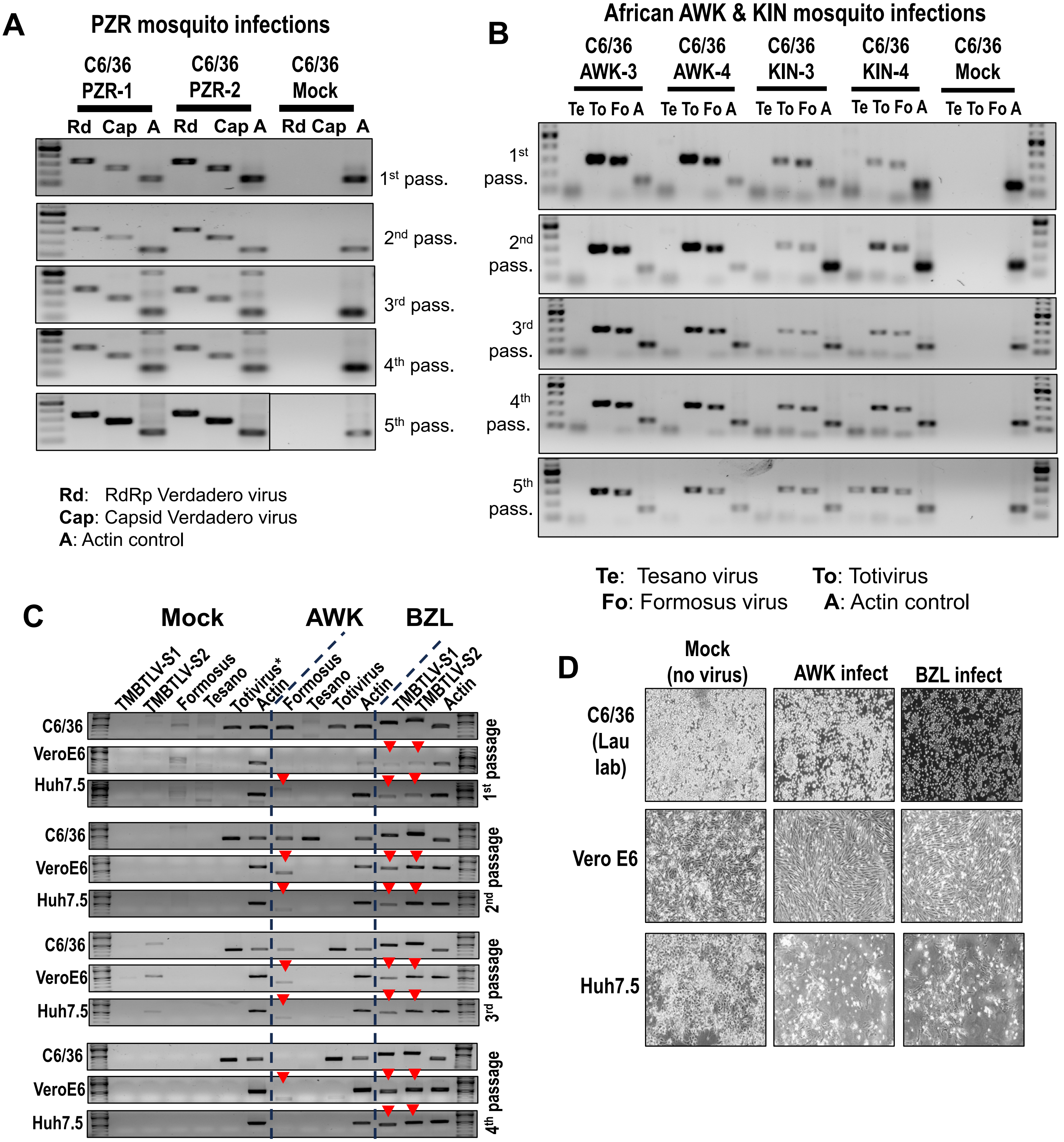


**Figure S5. Tracking virus presence with RT-PCR during blind passaging of C6/36 mosquito cells from infections with PZR, AWK and KIN mosquito homogenates – with the potential to infect mammalian cells.**
(A) RT-PCR of viral amplicons against the Verdadero virus RdRP and capsid genes. Two independent infections with two batches of PZR mosquitoes. (B) RT-PCR of viral amplicons against Tesano virus, Totivirus and Formosus virus from two independent infections with two batches of AWK and KIN mosquitoes. The actin control are primers against *Ae. albopictus* actin. (C) Continued blind passaging of virus stocks from C6/36 cells are then tested for infection of mammalian VeroE6 and Huh7.5 cells followed by RT-PCR detection of viral amplicons. Red arrowheads point to insect virus amplicons detected in the mammalian cells. Asterisk marks a totivirus already present in this lab stock of C636 cells. (D) Brightfield images cells from after 1 week of infection from 4th passage in (C) before harvesting total RNA for RT-PCR analysis. The cytopathic effect of infection from the BZL virus stock is most evident on C6/36 and Huh7.5 cells.
