## Supplemental Figure S6 for "Global genomics of *Aedes aegypti* unveils widespread and novel infectious viruses capable of triggering a small RNA response"

**Gupta, Sharma, et al.**
**Figure S6.**

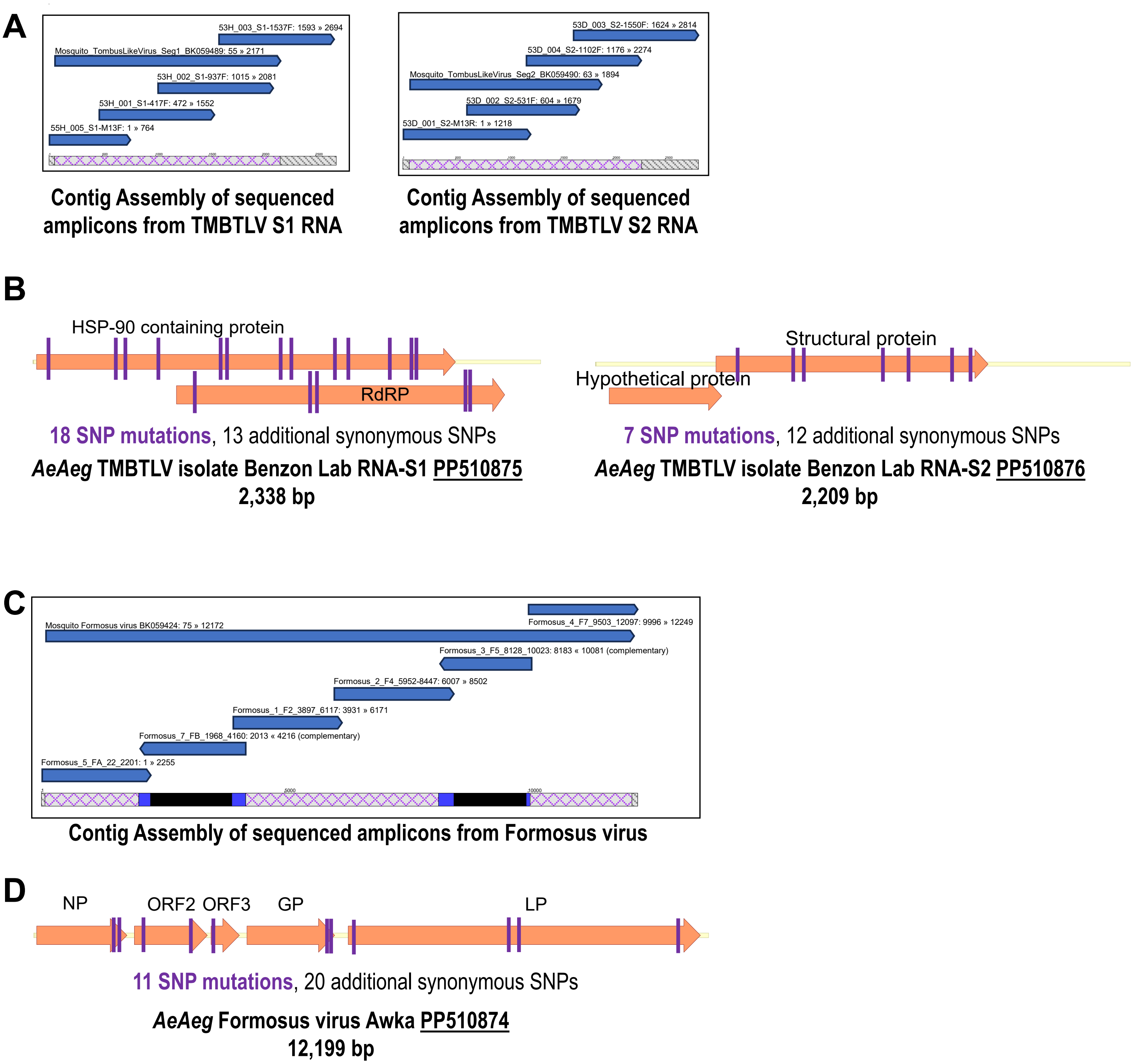


**Figure S6. Cloning and sequencing the TMBTLV and Formosus virus from *Ae. aegypti*.**
(A) Contig assembly of Sanger sequencing of cloned TMBTLV amplicons from S1 and S2 RNAs. (B) Diagrams of SNP mutations from our cloned fragment sequencing versus the initial TMBTLV references BK059489 and BK059490. (C) Contig assembly of Nanopore sequencing of cloned Formosus virus amplicons. (D) Diagrams of SNP mutations from our cloned fragment sequencing versus the initial Formosus references BK059424. Both of our TMBTLV and Formosus virus variants sequences have now been contributed to GenBank with verified sequence accessions that should facilitate its inclusion in the future updates of the GBVRL database.
